# Microbial Growth in an Enceladus Ocean Analog Medium Informed by Mineral Stability Modeling

**DOI:** 10.64898/2026.06.29.735333

**Authors:** Sabrina M. Elkassas, Tucker Ely, Teodora Zhivkova, Amina Patterson, Katelyn Weeks, Vinitra Nathan, Lydia Hayes, Sarina Mitchell, Margrethe Serres, Everett Shock, Peter Girguis, Christopher German, Frieder Klein, Jeffrey Seewald, Julie A. Huber

## Abstract

Evidence from the Cassini mission confirmed that Saturn’s moon Enceladus hosts a subsurface alkaline ocean where rock-water reactions may generate redox disequilibria capable of supporting microbial metabolisms. To investigate potential microbial survival under simulated Enceladus ocean conditions, we used thermodynamic modeling to develop a salt formulation consistent with one possible Enceladus ocean composition and supplemented it with putative microbial energy sources to create a growth medium. The medium was inoculated with samples from diverse ocean world analog environments on Earth to determine which microorganisms could persist under Enceladus-like conditions. The microorganisms persisting in this geochemically bounded medium were heterotrophic, metabolically versatile bacteria with low carbon requirements. Genomic and physiological analyses further showed the presence of multiple stress-response pathways, sodium-based bioenergetic systems, osmoregulation strategies, and other adaptations consistent with survival in alkaline, low-nutrient settings. These results suggest that some stress-tolerant heterotrophic bacteria may serve as useful model organisms for life in Enceladus’ subsurface ocean. These findings demonstrate the value of geochemically modeled media as a framework for constraining habitability, identifying relevant biosignatures, and probing potential microbial survival strategies beyond Earth.

## 1. INTRODUCTION

Ocean-bearing moons in our solar system represent some of the most compelling environments for astrobiological investigation, especially those where the saltwater oceans are understood to be in contact with a rocky interior, offering a key mechanism to providing a habitable environment that could, potentially, sustain life. Prior to the Cassini-Huygens (1997-2017), evidence for subsurface oceans on icy moons was derived primarily from telescopic observations and remote sensing measurements, including observations from the Pioneer, Voyager, and Galileo spacecraft missions (reviewed in Dougherty et al., 2018). However, in July 2005, Cassini’s flyby of Saturn’s moon Enceladus provided unexpected evidence for jets of water streaming from the moon’s south pole, inferred to originate directly from its ocean (Porco et al., 2006). Although it had been hypothesized decades earlier that plumes from Enceladus were the source of Saturn’s E-ring (Pang et al., 1984; Dougherty et al., 2018), Cassini’s direct sampling confirmed this connection and revealed the plumes contain salts, organic matter, and other compounds consistent with geochemical conditions that could potentially support life within Enceladus’ global ocean (Spahn et al., 2006; Postberg et al., 2009, 2011; Waite et al., 2006, 2009).

Subsequent Cassini flybys prompted a surge of studies interpreting spectral data from the Visual and Infrared Mapping Spectrometer (VIMS) and compositional measurements from the Cosmic Dust Analyzer (CDA) to infer the chemical and physical properties of Enceladus’ subsurface ocean. Silica nanoparticles from Saturn’s E-ring and sodium-rich ice grains in Enceladus’ plume have been interpreted to result from ongoing hydrothermal activity and water-rock reactions involving reduced mineral phases (Hsu et al., 2015). Modeling of Cassini data further suggests an alkaline to hyperalkaline ocean (pH 8 to ∼12.5; Zolotov, 2007; Postberg et al., 2009, 2011; Glein et al., 2015; Hsu et al., 2015; Glein et al., 2018; Fifer et al., 2022; Glein et al., 2025) enriched in molecular hydrogen (H_2_; Waite et al., 2017) and complex organic compounds (Postberg et al., 2018; Khawaja et al., 2025). Disequilibria between H_2_ and carbonate species in the ocean as described by Waite et al. (2017) implies the availability of metabolic energy for chemolithoautotrophic processes, such as methanogenesis. Hydrothermal temperatures exceeding 50°C inferred for Enceladus’ subseafloor environments (Sekine et al., 2015; Hsu et al., 2015) are consistent with fluid-rock reactions (Daval et al., 2022; Glein et al., 2020) that generate reduced gases and redox-active species capable of sustaining chemosynthetic life. Finally, more recent analyses of plume particles captured by Cassini have demonstrated the presence of sodium phosphates, confirming that key bioessential elements are available within the ocean (Postberg et al., 2023; Randolph-Flagg et al., 2023).

Motivated by this evidence for hydrothermal activity, available phosphate and energy for microbial metabolism, organics, and temperatures compatible with life, we used geochemical modeling to design an experimental microbial growth medium that reflected a predicted Enceladus seawater compositional space. Rather than to target specific microbial metabolisms *a priori*, the design of the Enceladus medium was guided by a thermodynamic modeling framework intended to constrain fluid composition based on plausible water-rock reactions and bulk geochemical observations. However, some assumptions about temperature, pressure, pH, major ion composition, and the inclusion of dissolved inorganic carbon and trace organics, were necessary to bound the model output by defining the chemical space within which equilibrium and speciation calculations were performed. Within these constraints, the model was used to predict the relative availability and speciation of dissolved components, which informed the final salt composition of the medium. Thus, while the model itself did not explicitly select for particular microbial metabolisms, the assumptions used to parameterize it reflect informed decisions about Enceladus-relevant conditions and indirectly shape the metabolic opportunities available in the resulting medium.

Whereas most geomicrobiological media are intentionally designed to enrich for specific metabolic groups, our approach uses geochemical modeling to infer ocean composition independent of metabolic considerations, allowing habitability to be evaluated while minimizing presumptions about the metabolic repertoires of microbes. We then inoculated the medium with samples from a range of Earth-based ocean world analog environments (Stern et al., 2025), including deep-sea hydrothermal vents, serpentinite mud volcanoes, and terrestrial basaltic lava caves. These efforts expand the discovery space for Earth’s microbial diversity and provide a systematic approach for evaluating possible physiological strategies and biosignatures of life in ocean worlds beyond Earth.

## 2. MATERIALS AND METHODS

### 2.1 Modeling

Our goal was to estimate a single ocean composition for Enceladus from a population of plausible chemical compositions consistent with the relatively few constraints available. Our population contains 50,000 independent fluid simulations, built by randomly selecting the total molal concentration of Fe, Mg, Ca, K, Sr, S, Br, F, B, C, Si, P and N from between 10^-12^ and 10^-1^, and Si, Cl and Na from between 10^-4^ and 10^-0.5^. pH and log *f*O_2_ were similarly drawn from broad distributions (6 → 12 and -20 → -1, respectively). This pH selection was designed to encompass the current estimates of fluid pH on Enceladus (pH 8 to ∼12.5; Zolotov, 2007; Postberg et al., 2009, 2011; Glein et al., 2015; Hsu et al., 2015; Glein et al., 2018; Fifer et al., 2022; Glein et al., 2025), and a large range of *f*O_2_, with the most oxidized systems (log *f*O_2_ ∼ -1) consistent with Earth’s seawater. Charge balance was achieved in each initial fluid by allowing either Na or Cl (randomly assigned) to be adjusted. This brought systems into charge balance by affecting the distribution of charge-bearing species, such as Na^+^ and Cl^-^, and their complexes.

Each initial fluid composition was brought to equilibrium with magnesite (MgCO_3_) through titration, allowed to speciate, and allowed to reach thermodynamic equilibrium with other saturated mineral phases. This converted our initial population of fluids into a population consistent with the presence of the mineral magnesite, because it aligns with interpretations of Enceladus’ plume chemistry (Glein and Waite, 2020) and remains broadly compatible with seawater compositions. This approach was also chosen for computational efficiency: forcing magnesite ensured that all simulations converged on magnesite-buffered solutions, rather than relying on rejection sampling to identify the subset of fluids that naturally achieved saturation. Mineral forcing therefore both accelerated the computation and constrained the fluid space by limiting degrees of freedom through mineral precipitation.

Our approach produced a range of seawater compositions that collectively explore a large region of the available parameter space but are all consistent with the equilibrium presence of magnesite. Moreover, carbon remained largely in C^4+^-bearing species. Sulfur was allowed to speciate among sulfate and sulfide bearing species depending on the final *f*O_2_ in each model. The final range of pH and *f*O_2_ values was affected both by the input values, and by the secondary precipitates which formed. Each individual seawater simulation was conducted with EQ3/6 Version 8 (Wolery & Jarek, 2003), using the methods and thermodynamic data described in Ely et al. (2023). The collective population of seawater compositions resulted in the precipitation of numerous other mineral phases. However, the most common were talc and quartz from carbonation, consistent with the predictions of Glein and Waite (2020; **Figure 1**).

**Figure 1.**
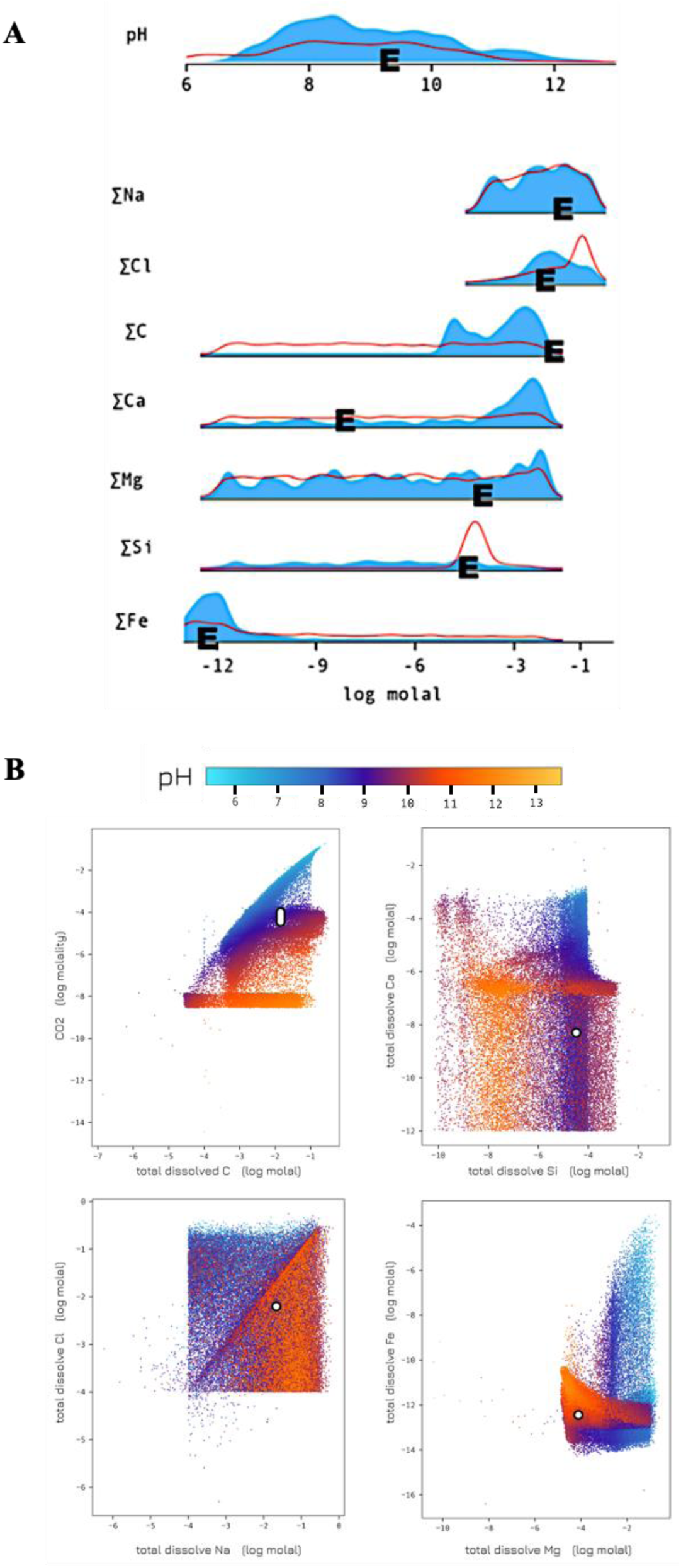
**A.** The distribution in concentrations of select elements, with “E” representing the average of the populations generated from the two modeled chemical equilibrium systems (magnesite alone shown in blue and magnesite-talc-quartz-ss shown in red, with peaks representing where simulations are clustered) at pH 9.2. The element labels on the left refer to the total dissolved abundance of these elements in solution among the population of models. Each element is distributed across several dissolved salt species (not shown). “E” is the single system that is represented by the elemental composition in **Table 1A**. For example, for total dissolved Si, seawater compositions in equilibrium with magnesite as the only mineral present can have nearly any dissolved Si concentration (the blue distribution is flat). In contrast, magnesite-talc-quartz-ss buffers an aqueous solution to a narrow range of dissolved Si concentrations (red peak). Other dissolved elements (not shown) were unaffected by mineral equilibration, displaying flat pre- and post-equilibration distributions between 10^-12^ and 10^-1^. **B.** Select dimensions of the full simulation output. White circles indicate the medoid composition before medium modifications (see **Materials and Methods** and **Table 2**). The molality of CO_2_ is restricted to a narrow range, as the medoid was calculated in dimensions of total element abundance, and not the dissolved species. Simulations cluster where individual fluids precipitate the same alteration minerals (talc and quartz).

To derive a single fluid composition from our population to be employed in the growth experiments, we calculated cluster-centers within the population. Our calculations resulted in five medoids^1^, of which we hand selected that with the largest pH. Medoids offer a way to convert a population of systems into one ‘most representative’ system as is necessary for experimentation (**Figure 1**). The choice of medoid number (how many clusters to subdivide a large population of points within), is a tradeoff between computational time and diminishing returns. Ultimately, we settled on five, which were selected to encompass a distribution in pH and chemical composition ‘clusters’, recognizing that the data population was heterogeneously clustered and therefore unfit for a single medoid among the total dissolved element dimensions and pH.

Our final simulated seawater composition was chosen as the experimental target (‘E’ in **Figure 1A**, in order of decreasing total concentration). ‘E’ locations don’t always align with the peaks for each element (e.g. C, Ca) is because the medoid calculation is trying to find the common best fit for all of the parameters described here, all of which co-vary together in n-dimensional space and are not independent of each other (**Figure 1B**). ‘E’ is dominated by total dissolved Na, C, and Cl (∼10s → ∼100s mmol), contains a moderate amount of Br, Mg, Si and S (∼10s → ∼100s µmol), and contains low concentrations of Ca, K, Sr, and Fe (< µmol) and is a combination of magnesite-only and magnesite-quartz-talc-solution series (ss) equilibrated model output. Our conditions are consistent with results from Glein and Waite’s (2020) model, with serpentinization as the primary water-rock reaction controlling Enceladus’ ocean chemistry, and carbonation occurring as a secondary process that buffers dissolved inorganic carbon and pH under alkaline conditions.

The elemental composition of the simulated seawater is shown in **Table 1A**, while the associated recipe for the experimental salt composition is shown in **Table 1B**. We chose to present this composition in terms of the total dissolved abundance of each element instead of aqueous species, because the solution is constructed in the lab by way of salt addition to 1 kg of pure water. The pH of the model output was 9.2, with modifications to the medium outside of model predictions (acetate), included in the table to balance the amounts of the elements initially predicted by the model. This required adjusting Na concentration by decreasing the NaCl concentration. It is important to note that the output of the model does not dictate microbial energy sources; it only outputs the salt composition of the basal microbial medium.

**Table 1.** **A.** Model output for the magnesite and magnesite-talc-quartz-ss. All concentrations refer to the log molality of the total dissolved element listed in column 1. The target column represents the medoid returned from the seawater simulations, while the final column represents the actual composition consistent with the salt recipe. **B.** The salt composition also includes later amendments to the medium, such as acetate, to keep elemental compositions as close to the model predictions as possible. These are not exact, because we are using a limited number of salts, each with its own fixed stoichiometry of elements, to construct an experimental solution that does not reside exactly on an intersection of salt combinations. Bolded values in the bottom represent the composition used in the final microbial medium detailed in Table 2.

| A | Elemental Composition | Magnesite |  | Magnesite-Mg-talc-Quartz-ss |  |
| --- | --- | --- | --- | --- | --- |
|  | Element | Target (log molal) | Final (log molal) | Target (log molal) | Final (log molal) |
|  | Ca | -8.3087 | -8.3087 | -8.3087 | -8.3087 |
|  | Br | -3.9654 | -8.4139 | -3.9654 | -8.4139 |
|  | K | -8.4139 | -8.4139 | -8.4139 | -8.4139 |
|  | Fe | -12.4291 | -12.4291 | -12.4291 | -12.4291 |
|  | Cl | -2.2046 | -2.2678 | -2.0000 | -2.1095 |
|  | Mg | -4.0812 | -4.0812 | -4.0812 | -4.0812 |
|  | Na | -1.6722 | -1.7595 | -0.5000 | -0.5567 |
|  | C | -1.8746 | -1.9324 | -0.8000 | -0.7646 |
|  | Sr | -8.9238 | -8.9238 | -8.9238 | -8.9238 |
|  | S | -6.5776 | -6.5776 | -6.5776 | -6.5776 |
|  | Si | -4.4999 | -4.4999 | -4.4999 | -4.4999 |
|  | N | - | - | - | - |
| B | Salt Composition | Magnesite |  | Magnesite-Talc-Quartz-ss |  |
|  | Salt | mmolal | mg | mmolal | mg |
|  | Na<sub>2</sub>CO<sub>3</sub> | 0.377399 | 40.00000 | 97.185651 | **10300.58465** |
|  | NaHCO<sub>3</sub> | 0.238076 | 20.00000 | 61.303669 | **5149.931778** |
|  | NaCl | 1.010258 | 59.0419 | 7.605688 | **444.495179** |
|  | NaNO<sub>3</sub> | - | - | 7.502920 | **637.708237** |
|  | NaOH | 9.859946 | 394.3693 | - | - |
|  | NaCH<sub>3</sub>COO | 5.534104 | 453.9868 | 6.730559 | **552.137311** |
|  | NH<sub>4</sub>Cl | 5.067378 | 271.0599 | - | - |
|  | MgCl<sub>2</sub> • 6 H<sub>2</sub>O | 0.082673 | **16.80756** | 0.082673 | **16.80756** |
|  | SiO<sub>2</sub> | 0.031632 | **1.90061** | 0.031632 | **1.90061** |
|  | MgSO<sub>4</sub> • 7 H<sub>2</sub>O | 0.000264 | **0.065188** | 0.000264 | **0.065188** |
|  | CaCl<sub>2</sub> • H<sub>2</sub>O | 0.000005 | **0.000722** | 0.000005 | **0.000722** |
|  | SrCl<sub>2</sub> | 0.000001 | **0.000189** | 0.000001 | **0.000189** |
|  | FeCl<sub>2</sub> • 4 H<sub>2</sub>O | 0.000000 | 0.000000 | 0.000000 | 0.000000 |
|  | KBr | 0.000004 | **0.000459** | 0.000004 | **0.000459** |

### 2.2. Microbial culture medium and conditions

The bolded salt compositions from **Table 1B** were converted to grams and modified into a microbial medium (per L) that could be prepared in the lab, using standard anaerobic microbial cultivation practices (Nakagawa and Takai, 2008). Given evidence for fluid-rock-derived H_2_ and lack of detected dissolved O_2_ or major oxidants in plume material, Enceladus’ ocean is expected to be predominantly reducing and anaerobic (Waite et al., 2017; Glein and Waite, 2020). Unlike Europa, where radiolytic oxidants may be cycled into the subsurface ocean, oxidant delivery on Enceladus is believed to be minimal, further supporting anoxic ocean chemistry (Hand et al., 2009).

The Enceladus Ocean Microbial Medium (**Medium 042a**; **Table 2**) consists of (A) a basal salts medium (model output + acetate + Na-resazurin indicator), (B) a trace element solution (model output), and (C) post-autoclaving/purging additions (added H_2_/CO_2_ gas + NaOH + vitamin solution + Na_2_S • 9 H_2_O reducing agent). (A) contains 0.44 g of NaCl, 0.64 g of NaNO_3_, 0.55 g of NaCH_3_COO, 0.02 g of MgCl_2_ • 6 H_2_O, and 0.2 g of K_2_HPO_4_ per liter of MilliQ water, consistent with salt composition predicted from the model, with an added 150 µL Na-resazurin (0.2%) as a redox indicator. (B) contains: 0.7 g MgSO_4_ • 7 H_2_O (7.0 x 10^-5^ g/L), 0.008 g of CaCl_2_ • 2 H_2_O (7.5 x 10^-7^ g/L), 0.002 g SrCl_2_ (2.0 x 10^-7^ g/L), 0.001 g FeCl_2_ (1.0 x 10^-7^ g/L), and 0.005 g KCl (5.0 x 10^-7^ g/L) dissolved in 85 mL MilliQ water. To (B), 5.0 g NaSiO_3_ • 12 H_2_O dissolved in 15 mL 1.0 M KOH was added to bring the solution up to 100 mL volume.

**Table 2.**
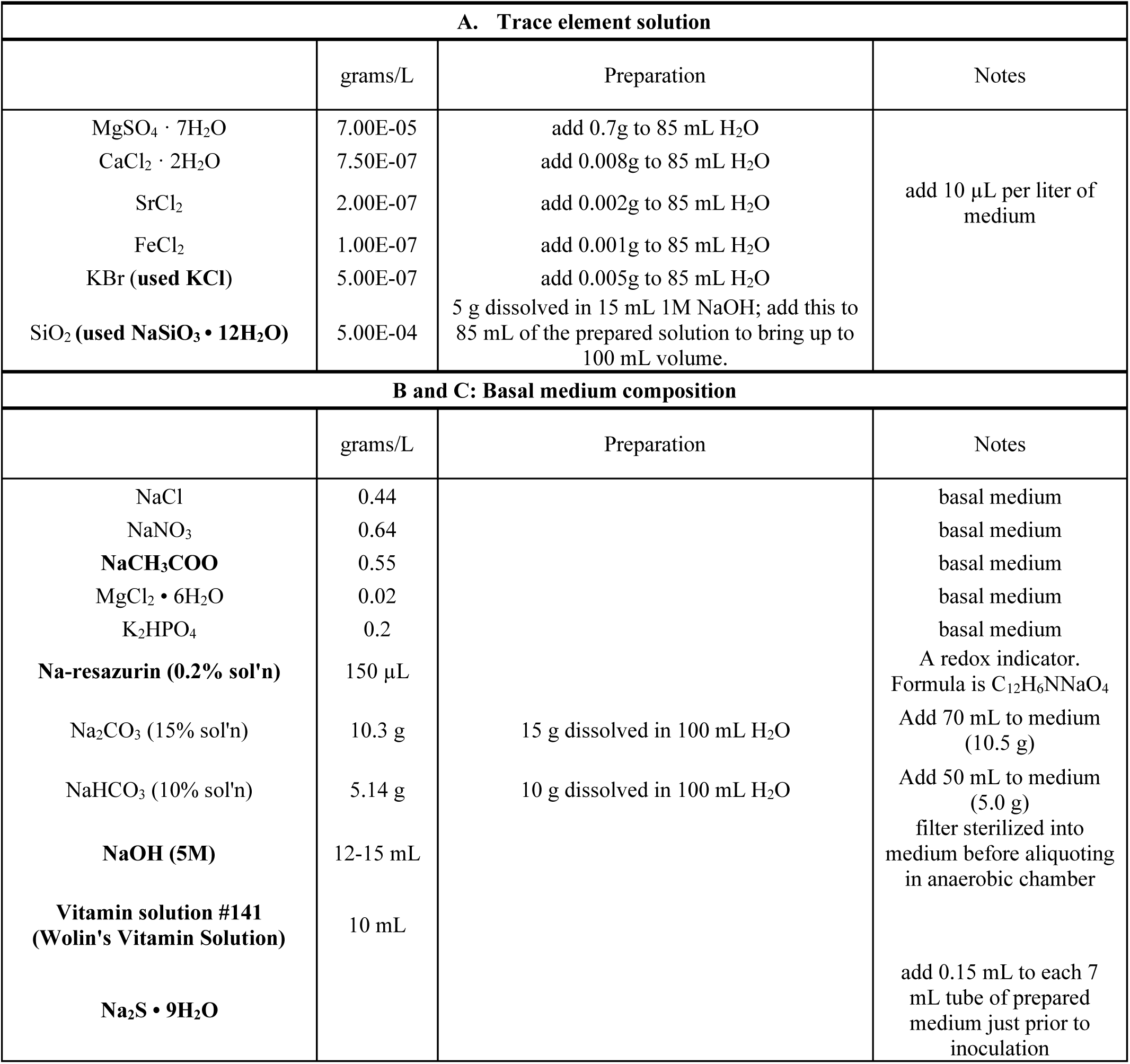
The Enceladus Ocean Microbial Medium 042a. Bolded constituents in column 1 represent additions/changes to the modeled salt composition.

To prepare Medium 042a, (A) was made in 855 mL of MilliQ water, and 10 µL of (B) was added. No additional trace element solutions were added. (A) was autoclaved then purged under a gas mix of 80/20 mol% H_2_/CO_2_ for 40 minutes or until cool. Culture vessels were over pressurized with the same H_2_/CO_2_ gas mixture. This mixture was based on the Enceladus plume composition (0.4-1.4 mol% H_2_ and 0.3-0.8 mol% CO_2_), predicted ongoing serpentinization reactions on Enceladus that generate H_2_, and the metabolic requirements of hydrogenotrophic methanogens, model organisms frequently studied in ocean worlds research (Waite et al., 2017; Seewald, 2017).

For (C), solutions of 15% (15 g/100 mL H_2_O) NaCO_3_, 10% (10 g/100 mL H_2_O) NaHCO_3_, Wolin’s Vitamin Solution (Wolin and Naylor, 1957) were filter-sterilized into autoclaved serum vials, sealed with a butyl rubber stopper and aluminum crimp, and made anoxic with 30 second N_2_ gas (100%) gas/vacuum cycles for 6 minutes total. Once (A) was cool and while still purging with 80/20 mol% H_2_/CO_2_, (C) was added: 70 mL of the 15% NaCO_3_ solution (10.5 g/L basal medium), 50 mL of the 10% NaHCO_3_ solution (5.0 g/L basal medium), followed by 10 mL of vitamin solution (another addition to the medium not predicted by the model) that was added slowly with a syringe fitted with a 0.2 µm syringe filter. The pH was adjusted to 11 (to match the highest predicted pH in Glein and Waite, 2020 and current estimates in Glein et al., 2025) using a pH probe with 10-15 mL of filter-sterilized, 5.0 M NaOH.

In an anaerobic chamber, 7 mL of the medium was dispensed into sterilized Hungate tubes, sealed, and, outside the chamber, purged using a benchtop gassing station in 30 second cycles with 80/20 mol% H_2_/CO_2_ gas/vacuum cycles for 3 minutes total per tube. Preparation details of the Medium 042a are in **Table S1**. A conceptual summary of the medium development is shown in **Figure 2**. During preparation of the pH 11 medium, no mineral precipitation occurred, which was unexpected given the challenges buffering high-pH medium (discussed in Sorokin, 2005).

**Figure 2.**
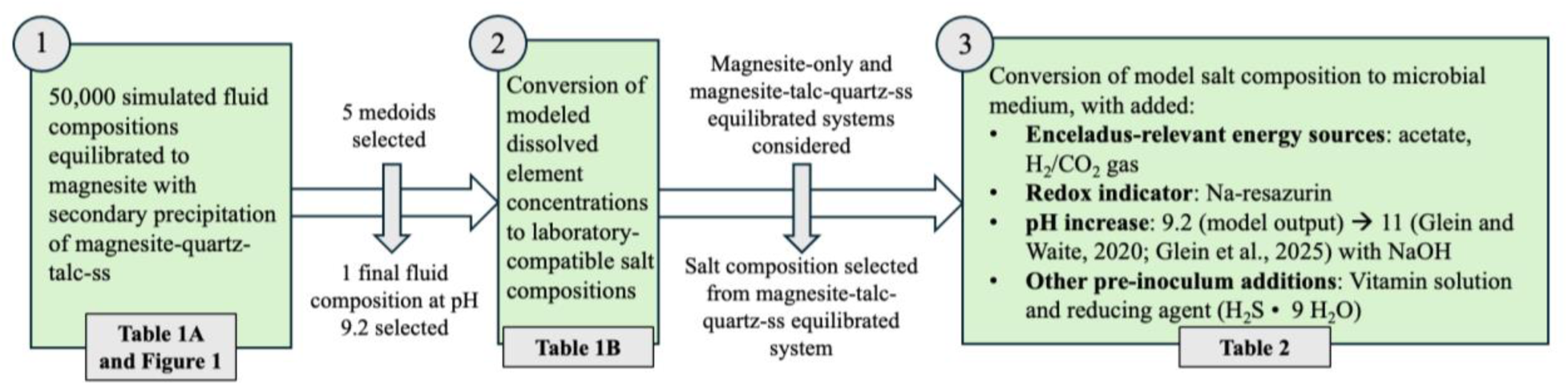
Conceptual summary of Medium 042a development.

### 2.3. Sample collection

Samples were collected at 5 field sites: **(#1)** serpentinite mud volcanoes at the Mariana forearc (Southwest Pacific Ocean), **(#2)** hydrothermal vents of the Mid-Atlantic Ridge, **(#3)** terrestrial hot springs (Oregon, USA), **(#4)** a bog atop the serpentinite section of the Josephine Ophiolite (Oregon, USA), and **(#5)** a basaltic cave (California, USA). Specifically, the serpentinized subseafloor fluids and sediment inocula (**#1**) were collected from four serpentinite mud volcanoes (Yinazao: 15.70949°N, 147.17665°E, 3666 meters below sea level (mbsl); Fantagnisña: 16.53758°N, 147.221035°E, 2019 mbsl; Asùt Tesoru: 18.110113°N, 147.101668°E, 1213 mbsl; South Chamorro Seamount: 13.784540°N, 146.002862°E, 2926 mbsl) along the Mariana forearc (Southwest Pacific Ocean) during cruise KM-2214 aboard the *R/V Kilo Moana* (November-December 2022). Fluid inoculum ranged in pH from 7.41 to 12.47. The sediment samples used as inoculum were collected from 12-14 cm depth below the seafloor. The geology and geochemistry from the serpentinite mud volcanoes have been extensively studied, reviewed in Fryer et al., 2020 and Eickenbusch et al., 2019. Diffuse hydrothermal fluids, microbial mats, and hydrothermal chimney material inocula **(#2)** were obtained during cruise FKT230303 aboard the *R/V Falkor(too)* (March-April 2023) from vent fields at the Kane Fracture Zone (23.66403844°N, 44.96518377°W, 2011 mbsl; Tucholke and Schouten, 1988), Grappe Deux (now Hydra; approximately 24.9°N, 45.5°W, 3791 mbsl; Baumberger et al., 2023), and Puy des Folles (20.5083°N, 45.6417°W, 1910 mbsl; Silantyev et al., 2024) on the Mid-Atlantic Ridge (Huber et al., 2024). Fluid inoculum (**#3**) was also collected from a small, bubbling hot spring in Oregon (Idleyld Park, Oregon; 43.293423°N, 122.365106°W; Mariner et al., 1990; collected with permission from the Umpqua National Forest) and a bog atop the serpentinized soil of the Josephine Ophiolite at Eight Dollar Mountain Botanical Area [**#4**; Selma, Oregon; 42.23320°N, 123.65900°W; Harper et al., 1984, collected under the Code of Federal Regulations, Title 43, Subtitle B, Chapter II, Subchapter C, Part 3800, Subpart 3809 (Surface use), subpart 3809.5]. Lastly, fluid inoculum (**#5**) was collected from a puddle on the floor filled with water dripping off a gold-colored biofilm on the cave ceiling in Catacombs Cave, Lava Beds National Monument (41.705°N, 121.515278°W; Taylor and Krejca, 2006). This puddle was about ∼800 feet from the cave entrance (far removed from sunlight) and harbored a surficial biofilm sitting atop the fluid. This sample was collected with permission from the Lava Beds National Monument.

Prior to inoculation, 0.15 mL 2.5% Na_2_S • 9 H_2_O under 100% N_2_ was added to reduce the medium for a final ΣH_2_S concentration of 0.05%. Inoculum size varied from 0.5-1 mL for fluid and sediment slurry samples. After inoculation, enrichment culture tubes were pressurized to 2 bar with 80/20 mol% H_2_/CO_2_ gas and incubated at 30°C. In total, 160 tubes of Medium 042a were inoculated from these field sites of opportunity. One culture was originally enriched in MJYTGL medium and was isolated after transferring to Medium 042a (isolate 8 described below).

### 2.4. Microbial isolation and DNA extraction

Growth of enrichment cultures was confirmed via microscopy. Enrichments showing signs of growth were transferred three times, and isolates were then obtained using dilution-to-extinction methods as described in Nakagawa and Takai (2008). This typically required 2 to 4 dilution series, until microorganisms were deemed axenic by checking morphological uniformity under the phase-contrast microscope (Zeiss Axioskop 40). Prior to extraction, duplicate or triplicate culture tubes of microbial isolate DNA were grown at 30°C until visibly turbid. One to two tubes of culture were used for DNA extraction, while the other was used for anaerobic cryogenic preservation in 100% glycerol, for a final concentration of 10% (1.8 mL culture + 0.2 mL 100% glycerol). DNA was extracted from 2-14 mL of culture using the Masterpure™ Complete DNA and RNA Purification Kit (LGC BioSearch Technologies) with added overnight DNA precipitation at -20°C. DNA was quantified using a Qubit™ 4 Fluorometer (ThermoFisher Scientific) with the 1x dsDNA High-Sensitivity Assay (Invitrogen).

### 2.5. 16S rRNA Sequence Preparation

DNA was extracted from 18 isolates, and the 16S rRNA gene was amplified, allowing for preliminary taxonomic identification of the isolates (**Table 3**, see **Supplementary Table 2** for an expanded table). Samples were prepared for paired-end 16S rRNA gene amplification using both 8F/1492R (Edwards et al., 1989; Stackebrandt and Liesack, 1993; and Galkiewicz et al., 2008) bacterial and 21F/958R (DeLong, 1989) archaeal primers. Polymerase chain reaction (PCR) mixtures for amplification of bacteria were composed of 27.8 µL nuclease-free water, 10 µL of 5x Green GoTaq Buffer (Promega) 5 µL of 10 µM 8F primer (Integrated DNA Technologies), 5 µL 10 µM 1492R primer (Integrated DNA Technologies), 1 µL of 10 mM dNTP mix (Thermofisher Scientific), and 0.2 µL GoTaq Polymerase (Promega) per reaction; for archaeal 16S amplification, the same mixture was used but with 5 µL each of 20 µM 21F and 958R primers (Integrated DNA Technologies). Each set of reactions was amplified for 30 cycles and visualized on a 1% agarose gel. Because archaeal DNA was not successfully amplified, only primers and PCR products for bacteria were submitted for 16S rRNA gene sequencing (Sequegen, Worchester, MA, USA).

**Table 3.** 16S rRNA gene identification of the 18 isolates.

| Isolate | Isolation Medium | Identifier | Site | 16S rRNA Sequence Identity, top hit, Genus and Species | 16S rRNA Sequence Percent Identity |
| --- | --- | --- | --- | --- | --- |
| 1 | Medium 042a | EB4 | Fantangisña mud volcano, Mariana forearc (Western Pacific Ocean) | *Marinobacter shengliensis* | 98.46% |
| 2* | Medium 042a + add'l acetate | 19B | Catacombs Cave (California) | *Hoeflea algicola* | 84.87% |
| 3 | Medium 042a + add'l acetate | 27B | Catacombs Cave (California) | *Paenisporosarcina indica* | 96.57% |
| 4* | Medium 042a + add'l acetate | 41B | Hot spring at Umpqua National Forest (Oregon) | *Marinobacter* sp. LQ44 | 97.63% |
| 5 | Medium 042a → Medium 042a pH 9.5 | 23B | Catacombs Cave (California) | *Paenisporosarcina quisquiliarum* | 96.94% |
| 6 | Medium 042a → Medium 042a pH 9.5 | 24B | Catacombs Cave (California) | *Paenisporosarcina indica* | 97.19% |
| 7 | Medium 042a → Medium 042a pH 9.5 | 26B | Catacombs Cave (California) | *Paenisporosarcina indica* | 96.81% |
| 8 | pH 10.5 MJYTGL → Medium 042a + add'l acetate | 18B | Hot spring at Umpqua National Forest (Oregon) | *Marinobacter vinifirmis* | 98.55% |
| 9 | Medium 042a → MJYTGL pH 10.5 | 48B | Hot spring at Umpqua National Forest (Oregon) | *Paraclostridium bifermentans* JCM 1386 | 98.49% |
| 10* | Medium 042a → MJYTGL pH 10.5 | 6B | Yinazao mud volcano, Mariana forearc (Western Pacific Ocean) | *Alishewanella agri* BL06 | 98.49% |
| 11 | Medium 042a → Marine agar #2216 (aerobic) | 21B | Catacombs Cave (California) | *Dietzia maris* | 95.66% |
| 12 | Medium 042a → Marine agar #2216 (aerobic) | 22B | Catacombs Cave (California) | *Paenisporosarcina indica* | 98.67% |
| 13 | Medium 042a1 → Marine agar #2216 (aerobic) | 28B | Catacombs Cave (California) | *Dietzia kunjamensis* | 94.21% |
| 14 | Medium 042a → 195c. + acetate | 36B | Catacombs Cave (California) | *Dietzia maris* | 98.03% |
| 15* | Medium 042a → 195c. | 40B | Hot spring at Umpqua National Forest (Oregon) | *Sulfurospirillum alkalitolerans* | 97.38% |
| 16 | Medium 042a → MJYTGL pH 8 | 34B | Catacombs Cave (California) | *Vibrio metschnikovii* | 99.03% |
| 17 | Medium 042a → MJYTGL pH 8 | 35B | Hot spring at Umpqua National Forest (Oregon) | *Ornithinibacillus caprae* | 95.20% |
| 18 | Medium 042a → Cami NO<sub>3</sub> | 33B | Hot spring at Umpqua National Forest (Oregon) | *Vibrio metschnikovii* | 95.53% |

\* = selected for whole genome sequencing

16S rRNA sequences were quality controlled, trimmed, and paired using DNA Subway (on CYVERSE; Hilgert et al., 2014). Consensus sequences were uploaded to nucleotide BLAST (nBLAST; Altschul et al., 1990) using the rRNA/ITS databases as reference and selecting sequences from only type material. Neighbor-joining phylogenetic trees (1000 bootstraps) and dissimilarity matrices were constructed using Molecular Evolutionary Genetics Analysis (MEGA v11; Tamura et al., 2021) with the 50 closest 16S rRNA BLAST hits, and edited using Interactive Tree of Life (iTOL, v6; Letunic and Bork, 2007) and Affinity Designer.

### 2.6. Whole Genome Sequencing

After examining taxonomy, neighbor-joining phylogenetic trees (**Figures 3** and **4)**, and dissimilarity matrices for all isolates, we chose to sequence the whole genomes of five microbial isolates, based on 16S rRNA gene dissimilarity to known isolates and whether we obtained the same microbial group from two different environments sampled. Samples were processed for genomic sequencing using methods described in Elkassas et al. (2024). This involved preparing sequence libraries using the Illumina DNA Prep, (M) tagmentation kit, and Integrated DNA Technologies for Illumina DNA/RNA Unique Dual Indexes per manufacturer protocols. Sequencing was performed on the Illumina NextSeq2000 platform using a 300-cycle flow cell kit to produce 2 × 150 bp paired-end reads at SeqCoast Genomics (Portsmouth, NH).

**Figure 3.**
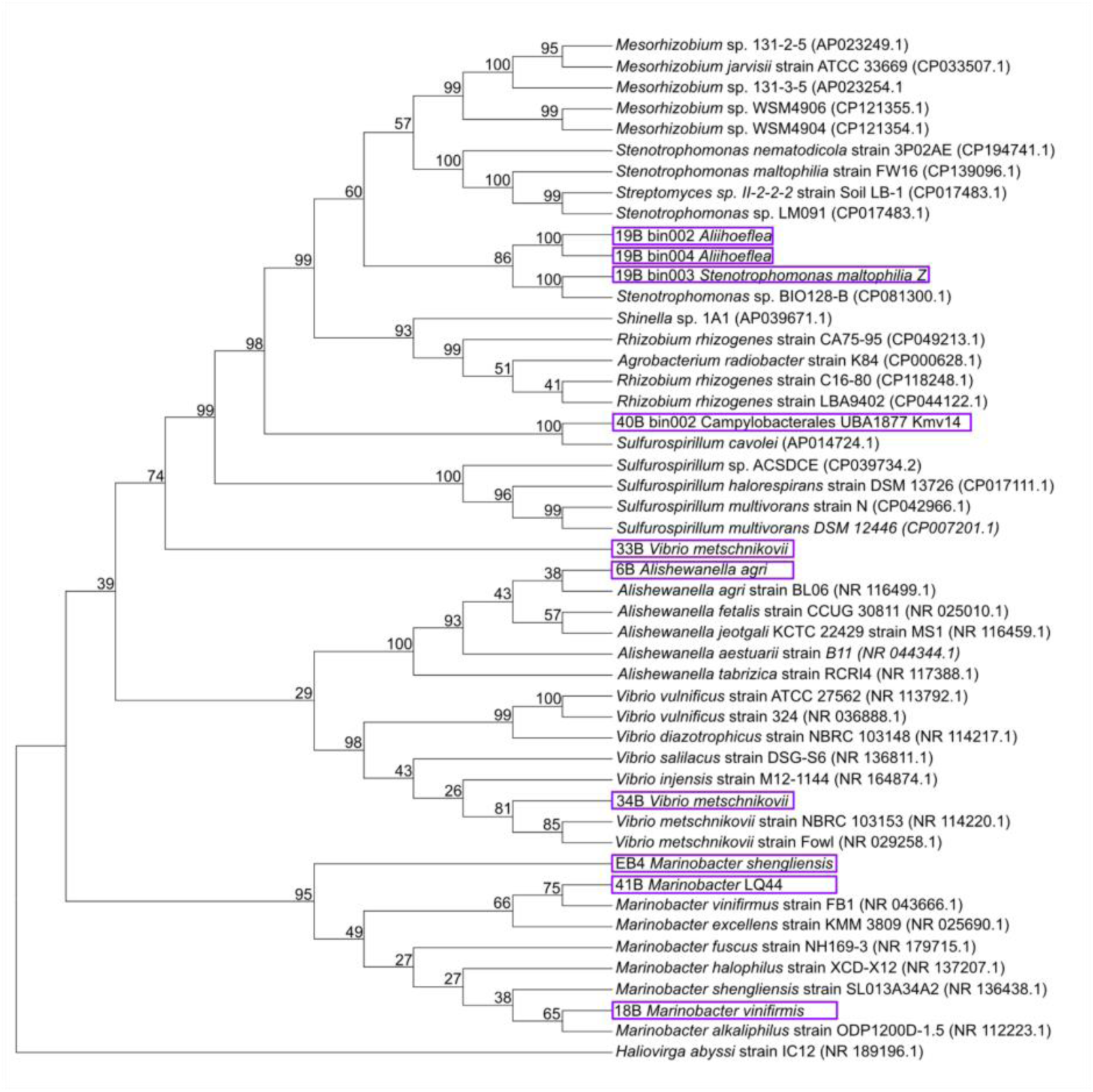
Neighbor-joining (Saitou and Nei, 1987) phylogenetic tree of all Proteobacterial isolates (indicated with a purple box) from initial enrichments in Medium 042a with the 5 closest related 16S rRNA sequences to each microbial isolate. The numbers preceding the isolates are the culture identifier. *Haliovirga abyssi* strain IC12 is included as an outgroup. The percentage of replicate trees in which the associated taxa clustered together in the bootstrap test (1000 replicates) are shown next to the branches (Felsenstein, 1985). The evolutionary distances were computed using the Maximum Composite Likelihood method (Tamura et al., 2004) and are in the units of the number of base substitutions per site. Codon positions included were 1st+2nd+3rd+Noncoding. All ambiguous positions were removed for each sequence pair (pairwise deletion). Evolutionary analyses were conducted in MEGA11 (Tamura et al., 2021; Stecher et al., 2020).

**Figure 4.**
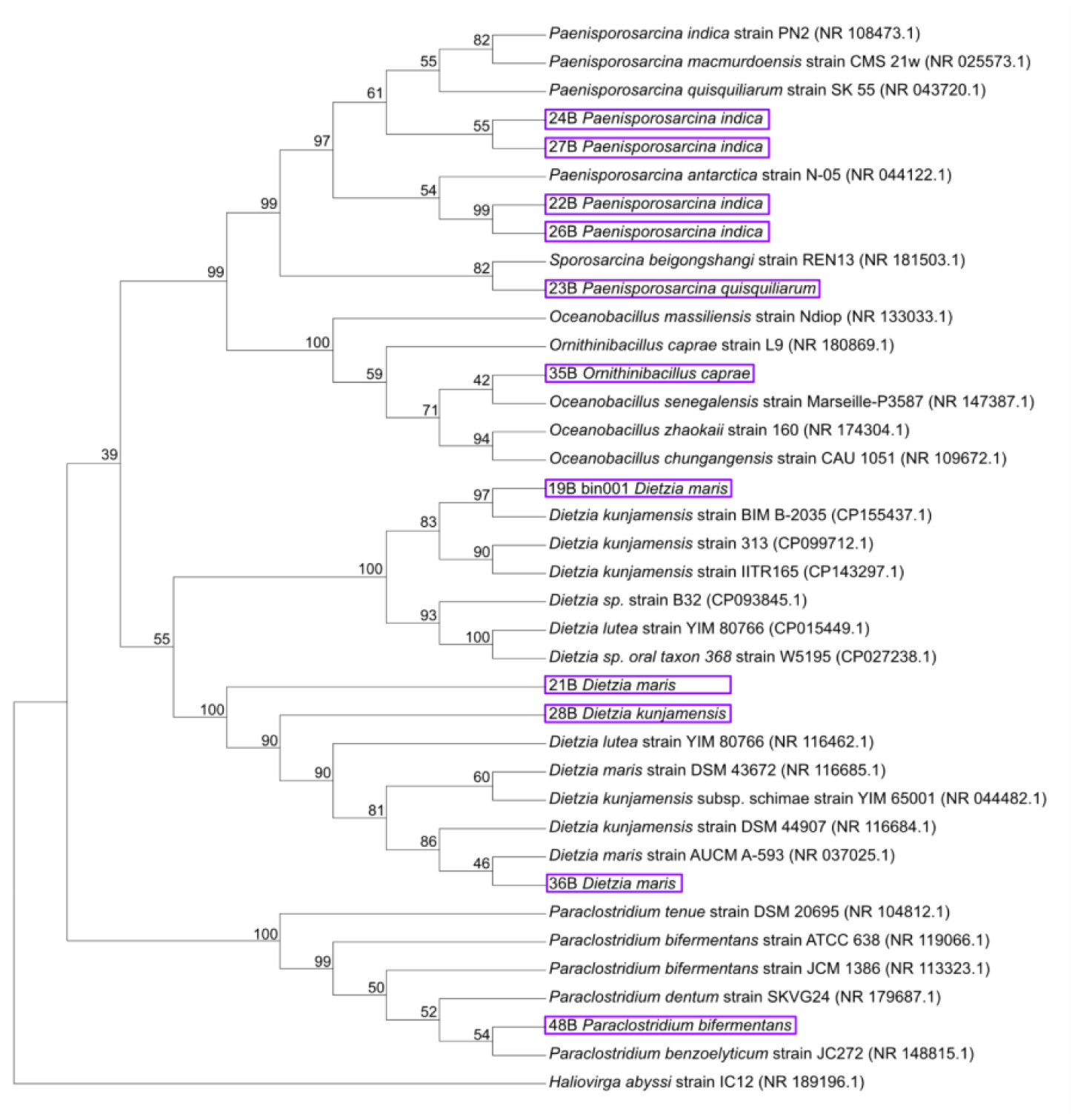
Neighbor-joining (Saitou and Nei, 1987) phylogenetic tree of all Bacillota and Actinomycetota isolates (indicated with a purple box) from initial enrichments in Medium 042a, with the 5 closest related 16S rRNA sequences to each microbial isolate. The numbers preceding the isolates are the culture identifier. *Haliovirga abyssi* strain IC12 is included as an outgroup. The percentage of replicate trees in which the associated taxa clustered together in the bootstrap test (1000 replicates) are shown next to the branches (Felsenstein, 1985). The evolutionary distances were computed using the Maximum Composite Likelihood method (Tamura et al., 2004) and are in the units of the number of base substitutions per site. Codon positions included were 1st+2nd+3rd+Noncoding. All ambiguous positions were removed for each sequence pair (pairwise deletion). Evolutionary analyses were conducted in MEGA11 (Tamura et al., 2021; Stecher et al., 2020).

### 2.7. Whole Genome Sequence Analysis

The whole genome sequences were analyzed using methods described in Elkassas et al. (2024) and uploaded to the Department of Energy Systems Biology Knowledgebase (KBase; Arkin et al., 2018). Sequences were quality-checked with FastQC v0.12.1 both before and after trimming (Lo and Chain, 2014), trimmed using Trimmomatic v0.36 (Bolger et al, 2014), assembled using SPAdes v3.15.3 (Bankevich et al., 2012), and checked for completeness and contamination using CheckM v1.0.18 (Parks et al., 2015).

Two of the five assembled genomes were determined by CheckM to be non-axenic, despite originating from cultures that appeared morphologically uniform, yielded only one 16S rRNA gene sequence from PCR-based screening, and had gone through multiple dilutions. These two assembled genomes were treated as metagenomes, and metagenome-assembled genomes (MAGs) were resolved from the genomes using a binning approach. Genomes were first indexed using BWA mem v7.17 (Li, 2013) and were then mapped to the indexed file. The resulting output was processed through Samtools, including converting .sam files to .bam files and sorting .bam files v1.13 (Li et al., 2009). The prepared sequences were binned using MetaBAT2 v2.15 (Kang et al., 2019), CONCOCT v1.1 (Alneberg et al., 2013), and MaxBin2 v2.2.4 (Wu et al., 2015) within KBase. An optimized bin set was selected from the binning outputs of all three binning software using DAS Tool v1.1.7 (Sieber et al., 2018). Bins were taxonomically classified using GTDB-tk v2.1.1 (Chaumeil et al., 2019). The bins were checked for completeness and contamination using CheckM v1.0.18 (Parks et al., 2015). All bins and the three pure culture genomes were quality-checked using Quast v4.4 (Gurevich et al, 2013).

Bins were filtered based on a completeness of >98% and contamination of <2%. Only bins that had >10% of reads recruited to them (using CoverM v0.7.0, Aroney et al., 2025) were kept for further analysis. The code for CoverM is available here: https://github.com/corporeal-snow-albatross-5/MAG-read-recruitment-using-CoverM. Quality-checked and filtered bins are hereafter referred to as metagenome-assembled genomes (MAGs). MAGs and the three pure culture genomes were taxonomically classified with GTDB-Tk v1.7.0 (Chaumeil et al 2019). Average nucleotide identities with all members of the same genus as the classified genomes were determined by FastANI v0.1.3 (Jain et al., 2018). Open reading frames predicted using Prodigal v2.6.3 (Hyatt et al, 2010) and annotated using DRAM v0.1.2 (Shaffer et al., 2020).

### 2.8. Pangenomic analysis

Pangenomic analysis of the MAGs and pure-culture genome sequences (8 total genomes) was conducted using the Anvi’o v8.0 (Eren et al., 2021) pangenome workflow (Delmont and Eren, 2018) to determine shared gene clusters among the 8 recovered genomes. First, assemblies generated from KBase were uploaded to Poseidon, the high-performance computing cluster at Woods Hole Oceanographic Institution. Using the Anvi’o contigs workflow (Eren et al., 2021), contig databases were made for each assembly, then sequence homology was determined (HMMER v3.4; Eddy, 2011), and open reading frames were predicted (Prodigal v2.1.1; Hyatt et al., 2010) and annotated with KEGGs, KOFAMs, and COGs. Gene cluster presence/absence was computed using the anvi-summarize function. The code used to generate the pangenome is located here: https://github.com/corporeal-snow-albatross-5/Enceladus_Medium_Pangenome.

We selected genes associated with anaplerotic inorganic carbon assimilation, pH homeostasis, temperature and osmotic stress tolerance, phosphate acquisition, oxidative stress defense, metal detoxification, aerobic respiration, and alternative anaerobic respiratory pathways, because these functions are broadly relevant to the physicochemical conditions inferred for Enceladus (see Glein et al., 2015; Waite et al., 2017; Postberg et al., 2018; Postberg et al., 2023; Randolph-Flagg et al., 2023) and are important adaptations for organisms surviving in analog environments on Earth (Merino et al., 2019). As such, the screened genes were selected not as unique biomarkers of life on Enceladus, but as genomic indicators of metabolic and stress-response strategies that could support persistence in an alkaline environment, with carbonate-dominated carbon chemistry, osmotic stress, and fluctuating redox conditions, as well as tolerance to reactive oxygen species and trace metals mobilized during hydrothermal alteration.

## 3. RESULTS

The designed microbial enrichment medium was named “Medium 042a”: “42” referencing *The Hitchhiker’s Guide to the Galaxy* (Adams, 1979). Medium pH was 9.5-10 prior to adding NaOH to raise the pH to 11 (Glein et al., 2025) and no mineral precipitation was observed. Out of 160 inoculations with environmental samples, we observed microbial growth in approximately half of these enrichment attempts, and all field sites (#1-5 in **Materials and Methods**) had at least one initial positive enrichment. There was growth in most initial enrichments for up to two transfers in Medium 042a, but many were unable to be propagated after these initial transfers without amendments. In some cases, several cultures that were enriched in Medium 042a were only successfully transferred and later purified to isolation in Medium 042a amended with additional acetate. In other cases, enrichments were transferred to a heterotrophic medium.

In total, seven microbial isolates were obtained solely through transfer and isolation using dilution-to-extinction in Medium 042a, and one was obtained from transferring an enrichment obtained in the heterotrophic, alkaline MJYTGL growth medium (Takai *et al*., 2005) into Medium 042a. Ten more isolates were obtained through transferring from Medium 042a to alkaline MJYTGL medium (Takai et al., 2005), circumneutral MJYTGL medium (Takai et al., 2005), circumneutral Marine Agar #2216 (Difco, Detroit, USA; ZoBell, 1941), and alkaline DSM 195c (for heterotrophic sulfate reducers; Alazard et al., 2003). In total, 18 bacteria were purified that were first enriched from the newly designed Medium 042a at pH 11, with 12 out of 18 isolated in high pH (>9.5) growth media (**Tables 3** and **S2**).

### 3.1. Taxonomy of microbial isolates

The taxonomic identities of all isolates based on 16S rRNA gene sequences or whole genome sequences when available, along with their sample origin, isolation medium, and average nucleotide identity (ANI), are summarized in **Tables 3** and **4**. All isolates were bacteria, spanning Phyla Pseudomonadota, Actinomycetota, Bacillota, and Campylobacteria (**Table 3**; **Figures 3** and **4**). Two of the isolates that returned a single 16S rRNA gene sequence resulted in mixed whole genome sequences, therefore metagenome-assembled genomes (MAGs) were binned to separate the specific taxa within the culture and annotate functions (see **Materials and Methods**; Isolates 2 and 15; **Table 4**). Based on both 16S rRNA gene and WGS, the isolates were identified as *Marinobacter shengliensis*, *Paenisporosarcina quisquiliarum, Paenisporosarcina indica* (four isolates), *Marinobacter vinifirmis* (two isolates), four MAGs from one culture: *Dietzia maris*, *Aliihoeflea* (two bins), *Stenotrophomonas maltophilia_Z, Alishewanella agri*, *Paraclostridium bifermentans*, *Ornithinibacillus caprae, Vibrio metschnikovii* (two isolates), *Dietzia maris* (two isolates), *Dietzia kunjamensis*, and one MAG from one culture, Campylobacterota UBA1877/Kmv14 (21 isolates total including MAGs, 18 cultures).

**Table 4.**
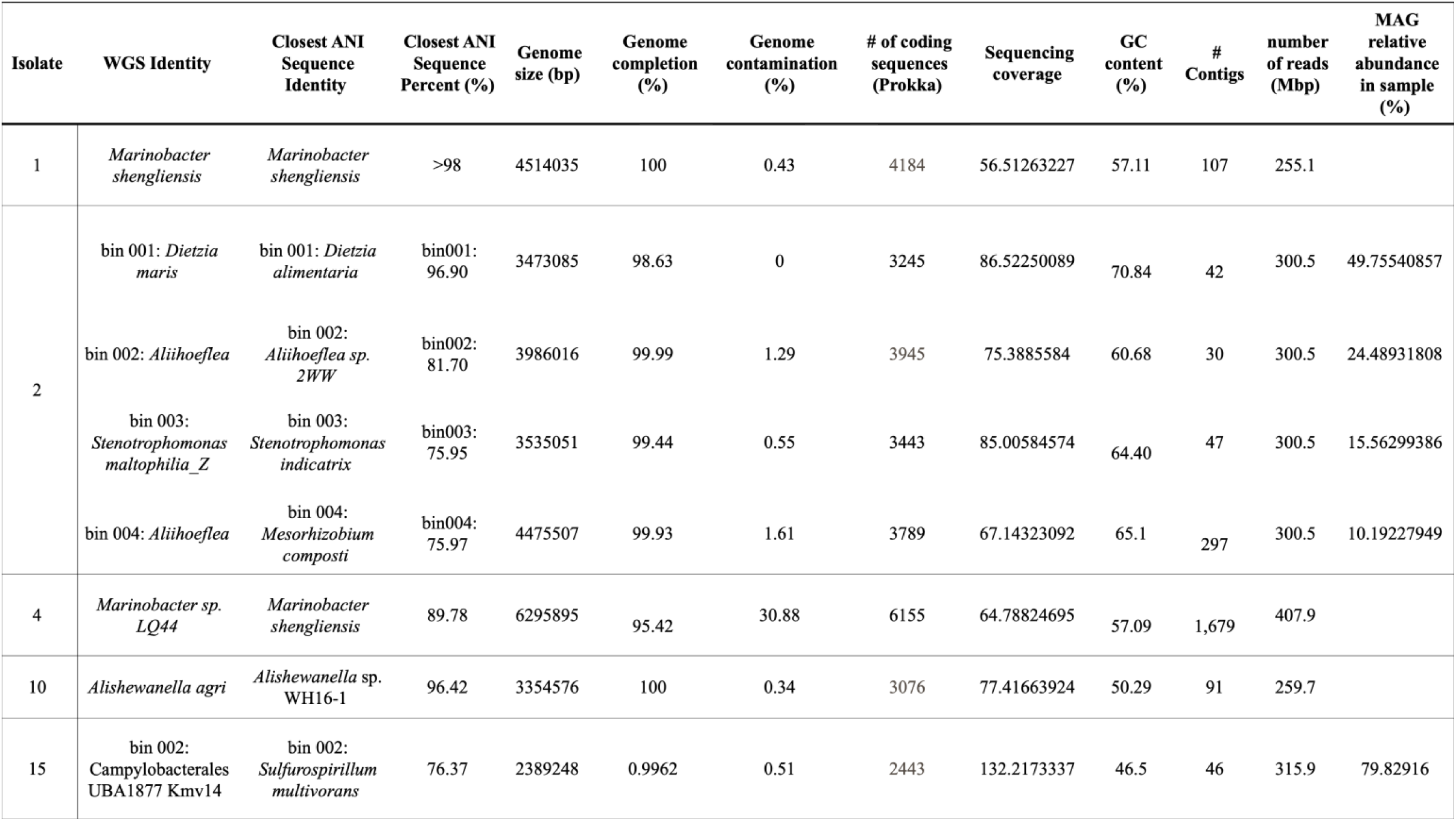
Identity and statistics of the five sequenced genomes.

| Isolate | WGS Identity | Closest ANI Sequence Identity | Closest ANI Sequence Percent (%) | Genome size (bp) | Genome completion (%) | Genome contamination (%) | # of coding sequences (Prokka) | Sequencing coverage | GC content (%) | # Contigs | number of reads (Mbp) | MAG relative abundance in sample (%) |
| --- | --- | --- | --- | --- | --- | --- | --- | --- | --- | --- | --- | --- |
| 1 | *Marinobacter shengliensis* | *Marinobacter shengliensis* | >98 | 4514035 | 100 | 0.43 | 4184 | 56.51263227 | 57.11 | 107 | 255.1 |  |
| 2 | bin 001: *Dietzia maris* | bin 001: *Dietzia alimentaria* | bin001: 96.90 | 3473085 | 98.63 | 0 | 3245 | 86.52250089 | 70.84 | 42 | 300.5 | 49.75540857 |
|  | bin 002: *Aliihoeflea* | bin 002: *Aliihoeflea sp. 2WW* | bin002: 81.70 | 3986016 | 99.99 | 1.29 | 3945 | 75.3885584 | 60.68 | 30 | 300.5 | 24.48931808 |
|  | bin 003: *Stenotrophomonas maltophilia_Z* | bin 003: *Stenotrophomonas indicatrix* | bin003: 75.95 | 3535051 | 99.44 | 0.55 | 3443 | 85.00584574 | 64.40 | 47 | 300.5 | 15.56299386 |
|  | bin 004: *Aliihoeflea* | bin 004: *Mesorhizobium composti* | bin004: 75.97 | 4475507 | 99.93 | 1.61 | 3789 | 67.14323092 | 65.1 | 297 | 300.5 | 10.19227949 |
| 4 | *Marinobacter sp. LQ44* | *Marinobacter shengliensis* | 89.78 | 6295895 | 95.42 | 30.88 | 6155 | 64.78824695 | 57.09 | 1,679 | 407.9 |  |
| 10 | *Alisewanella agri* | *Alisewanella sp. WH16-1* | 96.42 | 3354576 | 100 | 0.34 | 3076 | 77.41663924 | 50.29 | 91 | 259.7 |  |
| 15 | bin 002: *Campylobacteriales* UBA1877 KmV14 | bin 002: *Sulfurospirillum multivorans* | 76.37 | 2389248 | 0.9962 | 0.51 | 2443 | 132.2173337 | 46.5 | 46 | 315.9 | 79.82916 |

### 3.2. Genomic analysis of microbial isolates

A pangenomic approach was used to analyze the shared genomic content of the three microbial isolate genomes and five MAGs, focusing on carbon and energy metabolism, as well as adaptations to high pH and other environmental stressors such as temperature and oxidation stress (see **Materials and Methods**; **Tables S3** and **S4**, **Figure 5**). Genomic analysis of carbon metabolism revealed that all bacterial isolates and bins were heterotrophs capable of CO_2_ assimilation through anaplerotic pathways that replenish TCA cycle intermediates (Braun et al., 2021). Key enzymes involved in this process are PEP carboxylase, pyruvate carboxylase, malic enzyme, and carbonic anhydrase (**Figure 5**). The exception was *Dietzia maris* bin 001, which had both the lowest number of called marker gene sets and the lowest completion of all bins at 98.63%; **Table 4**). **None of the isolates had genes for carbon fixation** (Hügler and Sievert, 2011; Montoya et al., 2012). All isolates carried the genetic potential to utilize diverse carbon substrates, such as long-chain and aromatic hydrocarbons, sterols, and amino acids, along with acetate (**Table S5**).

**Figure 5.**
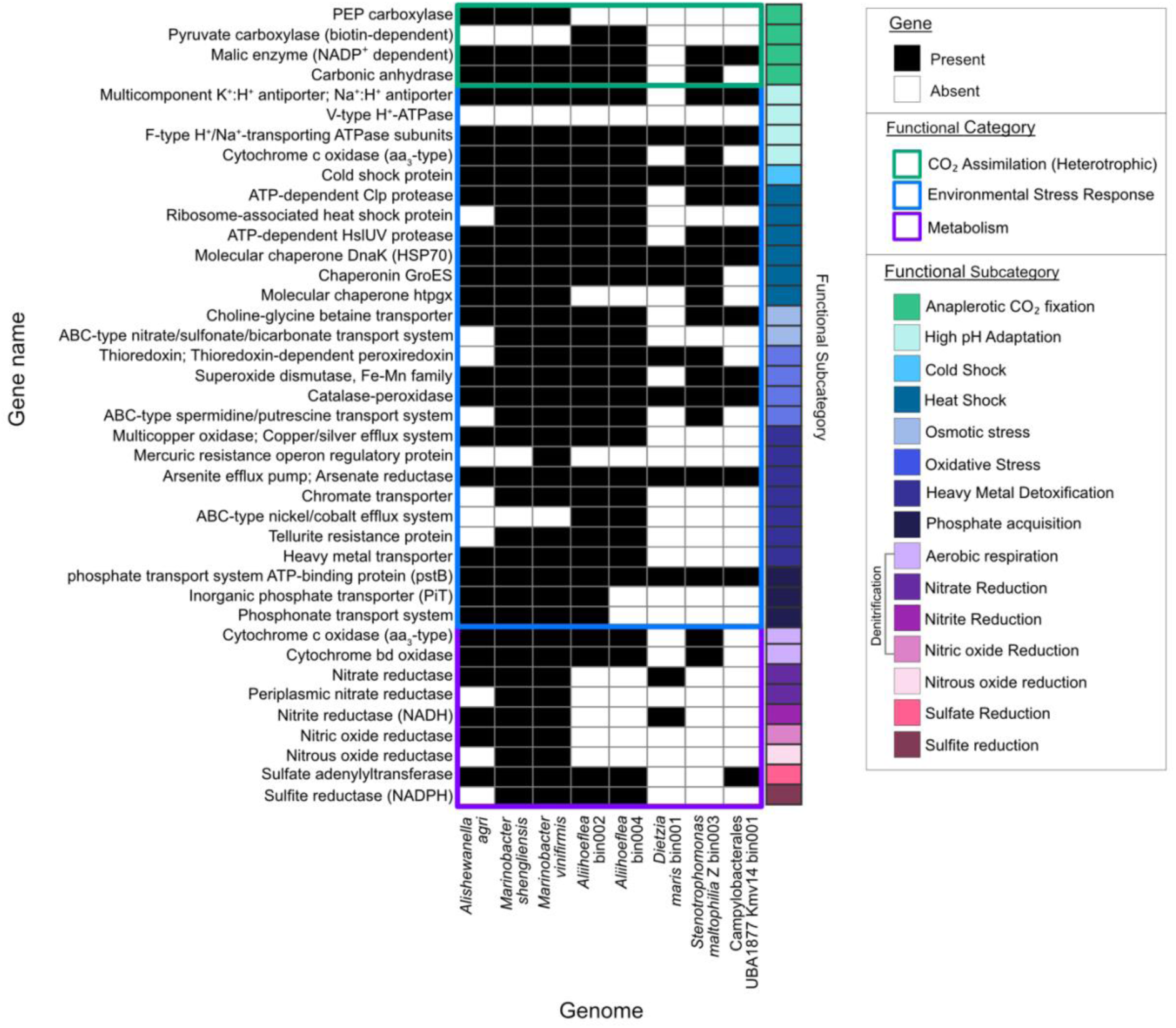
Presence/absence plot showing genes within sequenced genomes related to CO_2_ assimilation: anaplerotic CO_2_ fixation; environmental stress response: high pH adaption, cold shock, heat shock, osmotic stress, oxidative stress, heavy metal metabolism, phosphate acquisition; and metabolism: aerobic respiration, nitrate reduction, nitrite reduction, nitric oxide reduction, nitrous oxide reduction, sulfate reduction, and sulfite reduction.

With respect to dissimilatory metabolism, both *Marinobacter* isolates carried genes for complete denitrification, while *Alishewanella agri* and *Dietzia maris* bin 001 contained genes for partial denitrification. All isolates except for *Dietzia maris* bin 001 and *Stenotrophomonas* Z bin 003 had genes for sulfate reduction, and of these genomes, all but *Alishewanella agri* can also perform sulfite reduction. All but two bins (*Dietzia maris* bin 001 and Campylobacterales UBA1877 Kmv14 bin 001) also carried genes for aerobic respiration, indicating most isolates are facultative anaerobes. Notably, no genes for dissimilatory metabolism were shared among seven or more genomes.

All eight genomes contained genes for adaptation to environmental stressors, including F-type H^+^/Na^+^ ATPase for high pH adaptation, cold shock protein and molecular chaperone dnaK/heat shock protein 70 for thermal stress, catalase-peroxidase for oxidative stress, and the arsenite efflux pump/arsenate reductase for heavy metal detoxification. Genes encoding phosphate transport systems were detected across all eight genomes, including components of high-affinity ABC transporters (pstB) and low-affinity PiT transporters (pitA; four genomes). Seven of eight genomes (all except for *Dietzia maris* bin 001), contained genes for NADP^+^-dependent malic enzyme for replenishing tricarboxylic acid (TCA) cycle intermediates, multicomponent K^+^:H^+^ antiporter/Na^+^:H^+^ antiporter for high pH adaptation, heat shock proteins ATP-dependent Clp protease, HsIUV protease, and chaperonin GroES; osmotic stress response choline-glycine betaine transporter; and oxidative stress response protein superoxide dismutase, Fe-Mn family. For shared gene clusters across the genomes, see **Table S3**; for selected genes represented in **Figure 5**, see **Table S4**.

## 4. DISCUSSION

The goal of this work was to couple geochemical modeling of a possible composition of Enceladus’ subsurface ocean and translate it into a physical microbial enrichment medium to explore microbial physiology and adaptations from diverse ocean world analog environments on Earth to putative conditions in Enceladus’ ocean. The thermodynamic model constrained the basal salt composition, and because the dominant microbial metabolic strategies on Enceladus, if they exist, are completely unknown, the medium was not engineered to enrich for only one particular type of microbial metabolism. Instead, we modified the basal salt medium by removing O_2_ and adding H_2_/CO_2_ and acetate to keep the conditions consistent with Enceladus plume observations and modeling studies.

Our approach created an opportunity to identify physiological and genomic features associated with microbial growth at Enceladus-like conditions, thereby broadening the scope of exploration for life on both Earth and other ocean worlds. Of the isolates recovered from the designed Medium 042a, all genomes (including four MAGs) and 11 of the 12 isolates with only the 16S rRNA gene sequenced, likely represent novel species, based on <95% average nucleotide identity for genomes (ANI; Richter & Rosselló-Móra, 2009) and <98.7% 16S rRNA gene sequence identity (Stackebrandt and Ebers, 2006; Kim et al., 2014). The remaining isolates are likely novel strains, with >98.7% 16S rRNA gene sequence identity (Kim et al., 2014). The novelty of these isolates highlights previously unexplored microbial diversity capable of growth at high pH across diverse geological environments with growth medium informed by putative Enceladus ocean conditions.

The alkaline pH of the medium was perhaps the largest challenge for propagating the cultures to isolate a single microbial taxon. High pH conditions pose a substantial challenge for microbial cells, which must maintain an intracellular pH near 7 to 8 even when the surrounding environment contains an excess of hydroxide ions. This requires specialized physiological mechanisms such as selective substrate utilization and ion pumping systems (Dilworth and Glenn, 2007; Horikoshi, 2021). Consistent with this, the pangenome analysis showed that all genomes contained at least one gene involved in coping with alkaline stress, suggesting that these functions may be important for survival in Enceladus-like conditions, discussed further in **section 4.3**. Furthermore, six of the eighteen cultures were ultimately isolated in circumneutral pH medium, which suggests that even though all were initially enriched at pH 11, the lower pH conditions permitted more robust growth and initial enrichments were most likely at their growth limits for pH.

### 4.1. Culture transfer attenuation

From the initial enrichment cultures that exhibited growth in Medium 042a, many did not persist beyond two transfers without amendments or transferring to lower pH medium. Although vitamins were included in the medium, no trace elements or metals were added beyond those present in the model output (which contained trace amounts of Fe, Si, and Sr). With each transfer, any trace elements, metals, or other required nutrients from the original source sample were diluted by a factor of 8 (1 mL inoculum into 7 mL medium). Cultures that failed to propagate may have had trace element or metal requirement than could not be met after the second transfer. Cultivation of microorganisms, especially from extreme environments, is a notoriously challenging process (Alain and Querellou, 2009). Supplementing with a trace element solution, such as the widely used Wolin’s mineral solution (Wolin et al., 1963), might have promoted sustained growth of transfers, but without better constraints on Enceladus’ ocean chemistry, the chemical concentrations in existing formulations may not be appropriate. Instead, in the interest of cultivating strains for subsequent analyses, we relied on acetate amendments, transfer into a more organic-rich medium, or reduction of pH to below 8 to further enrich the isolates.

### 4.2. Uptake of organic carbon

There were two sources of organic molecules added to the Medium 042a: vitamin solution 141 (10 mL/L medium; Wolin and Naylor, 1957) and acetate (0.55 g/L medium) (**Tables 2** and **S1**). Vitamins are complex organic compounds required by autotrophs and heterotrophs alike that are used as cofactors or to build coenzymes; they are not typically used as a central carbon source (Koser, 1948). In the pangenomic analysis, isolates had broad biosynthetic capacity for core vitamins while exhibiting targeted dependencies on more complex cofactors. Folate (vitamin B9) biosynthesis genes were consistently present, including folD and upstream components such as folBCEKP, suggesting the capacity for extensive tetrahydrofolate-mediated one-carbon metabolism (Maden, 2000). The ability to process and transfer one-carbon units may be advantageous in high-pH environments, where inorganic carbon is dominated by carbonate species and dissolved CO_2_ is scarce (Glein et al., 2015). Similarly, pathways for riboflavin (vitamin B2) and pyridoxal phosphate (vitamin B6) synthesis were well represented, with genes including ribABDEH, and pdxAJ or pdxST, indicating metabolic independence for cofactors central to redox reactions and amino acid metabolism across diverse microbial taxa (Rodionov et al., 2002; Zhang and Gladyshev, 2008). In contrast, thiamine (vitamin B1) biosynthesis was incomplete in several genomes, with partial representation of thiCDEG, alongside transport systems such as thiBPQ, suggesting that some isolates rely on environmental thiamine or precursor salvage, a common strategy in nutrient-limited systems (Rodionov et al., 2002). The strongest dependency was observed for cobalamin (vitamin B12), as complete biosynthetic pathways involving cobAFILMNSTU were absent, while uptake systems including btuB and the btuCDF operon were detected, indicating reliance on exogenous sources or microbial cross-feeding (Seth and Taga, 2014). Most microorganisms in Earth’s oceans exhibit vitamin auxotrophy, especially with respect to vitamin B_12_ (cobalamin), which is metabolically complex and energetically expensive to produce (Sañudo-Wilhelmy et al. 2014; Raux et al., 2000). Isolates therefore appear to be largely self-sufficient for essential redox and C1-breakdown associated cofactors but depend on environmental or community-derived inputs for more complex vitamins, particularly cobalamin, supporting the amendment of vitamins to the medium.

Acetate, the second organic compound added to the medium, was not included in the initial model but was introduced as an amendment to the model based on the following logic. On Earth, the process of serpentinization produces H_2_ (Charlou et al., 2002; McCollom and Bach, 2009; Klein et al., 2013), which creates reducing conditions that can then favor the reduction of inorganic carbon to produce methane and other low-molecular-weight hydrocarbons (McCollom and Seewald, 2007), as observed in modern serpentinizing systems such as the Samail ophiolite (0.47-4.4 μM; Rempfert et al., 2017) and the serpentinite mud volcanoes of the Mariana forearc (1-42 µM; Eickenbusch et al., 2019). While the concentration of acetate in Enceladus’ ocean has not been quantified, the INMS and CDA instruments on Cassini detected organics up to ∼200 atomic mass units (acetate is ∼59 amu) in at least 4% of the ice grains able to be measured by CDA (Postberg et al., 2018). Analysis of fresh ice grains from the E5 high-speed Cassini flyby of Enceladus in Khawaja et al. (2025) indicate the tentative detection of aliphatic O-bearing species, including fragments consistent with acetic acid/acetate moieties, strengthening the case for active organic chemistry in the subsurface ocean. In addition, among water-soluble organics in carbonaceous chondrites, which compose the core of Enceladus, acetate is the most abundant (Yuen et al., 1984). Although vitamins were added to satisfy general biosynthetic requirements and acetate was included to mirror organics detected in Enceladus’ plume, rather than to serve as a primary carbon source, all isolates from Medium 042a (without transfer to another organic-rich medium) were heterotrophs and possessed the *acs* gene for acetate assimilation.

Given the minimal diversity and utility of the carbon sources added, we expected the microbial isolates purified from Medium 042a (three pure cultures total and five MAGs) to be highly selective carbon utilizers. However, we found that all the isolates carried genes with the potential to utilize diverse carbon sources, indicating they are likely heterotrophic generalists (see **Table S5**). Additionally, all but one isolate (the *Dietzia maris* bin with the lowest completion) also had genes for heterotrophic CO_2_ fixation, whereby CO_2_ is used to replenish TCA cycle intermediates, rather than fixed into sugars for biomass accumulation (Braun et al., 2021). While this strategy is energetically inefficient and unlikely to sustain growth in isolation, it is widespread among heterotrophic microorganisms in the deep ocean and other carbon-limited environments, where it can maintain metabolic flux when key intermediates are scarce (Manna et al., 2025). In such settings, CO_2_ assimilation replenishes TCA cycle intermediates such as oxaloacetate and malate in the absence of more complex organic inputs (Trembath-Reichert et al., 2021; Braun et al., 2021).

Anaplerotic CO_2_ assimilation does not provide sufficient carbon or energy to sustain continued biomass production, so it was likely insufficient to support prolonged growth without additional organic inputs, such as acetate, which was subsequently shown to stimulate growth. This interpretation is consistent with the absence of complete autotrophic carbon fixation pathways across all genomes, including the Wood-Ljungdahl pathway, Calvin cycle, and reductive TCA cycle (Hügler and Sievert, 2011; Montoya et al., 2012), indicating that these organisms are not capable of primary carbon fixation. Instead, they appear to rely on a metabolically flexible, predominantly heterotrophic lifestyle, supplemented by limited CO_2_ incorporation to buffer central metabolism under carbon-constrained conditions. Such a strategy may be particularly advantageous in environments analogous to Enceladus, where simple carbon compounds may be intermittently available, but more complex biosynthetic precursors are scarce.

### 4.3. Energy sources and stress response

Across isolates, there was no pattern for dissimilatory metabolic strategy, with some isolates carrying genes for growth under both aerobic and anerobic conditions, using nitrate, nitrite, or sulfite as alternative electron acceptors coupled to heterotrophy, whereas some were exclusively aerobic (a *Stenotrophomonas maltophilia Z* bin) or anaerobic (bins for *Dietzia maris* and Campylobacteriales UBA1877/Kmv14; **Figure 5**). Although they were isolated under anaerobic conditions, potential oxygen contamination could have occurred while field sampling if the original inoculum contained oxygen exceeding the capacity of the reducing agent (Na_2_S • 9H_2_O) or if no H_2_/CO_2_ gas was immediately available.

Given this diversity of metabolism, we focused our pangenomic analysis on stress response genes for a variety of environmental conditions, including high pH adaptation, cold and heat shock, osmotic stress, oxidative stress, and heavy metal detoxification (See **Materials and Methods**). The recovery of organisms not typically associated with high-pH environments, together with the consistent presence of stress response genes, suggests that persistence in our high pH, low organic, Enceladus-like medium selected for organisms with broad stress tolerance strategies, rather than strictly alkaliphily or the ability to survive at low organic carbon concentrations.

With respect to high pH adaptation, Na^+^:H^+^ and K^+^:H^+^ antiporters and F-type H^+^/Na^+^ ATPases, detected in most genomes recovered, are genes associated with alkaliphily, obligate growth above pH 9.0 (Horikoshi, 1999). Given our initial enrichment conditions in Medium 042a at pH 11, this adaptation is consistent with selection of obligate alkaliphiles. Despite that, some of the isolates were transferred to lower pH medium to propagate growth. A major challenge for organisms in alkaline environments is maintaining cytoplasmic pH, as plasma membranes become unstable above pH 9 (Horikoshi, 1999). In a study by Kitada and Horikoshi et al. (1978) with an alkaliphilic *Bacillus* spp., they found that when 0.2 N NaCl was added to the medium, the rate of incorporation of a test compound (α-aminoisobutyrate) was 20 times that with no NaCl amendment. This may mean the presence of available Na^+^ and other single-valent metal ions are determinants of survival in alkaliphilic organisms, providing a means to facilitate transport of H^+^ ions into the cell (Kitada and Horikoshi, 1978).

Another adaptation to high pH is the F-type H^+^/Na^+^ ATPase, a sodium-dependent, proton translocating ATP synthase that generates a chemiosmotic gradient using sodium-motive force to aid in ATP production when external proton concentration is low. In a study by Mulkidjanian et al. (2008) using structural and phylogenetic analyses, they found that utilization of a sodium gradient for ATP production is the evolutionary ancestor of the proton coupling ATP synthase used by most organisms today. These genetic and physiological adaptations help maintain intracellular pH between 7 and 8.5 in otherwise extreme conditions (Horikoshi, 1999). Since the F-type H^+^/Na^+^ ATPase was found in most genomes in this study and these enzymes are predecessors to modern ATP production machinery, we recommend looking into to the degradation products of these proteins and quantifying the amount and availability of monovalent metal ions in Enceladus’ ocean as a potential biosignature.

Another important stressor microbes face in the natural world is fluctuating changes in solute concentrations outside the cell, which can disrupt how water moves across the cell membrane, thus causing osmotic stress (Bremer and Krämer, 2019). Osmolytes, such as glycine betaine, proline, ectoine, and glutamate serve as a protective mechanism when microorganisms are under such osmotic stress, which can occur in a variety of environments such as hypersaline lakes and to a lesser degree, in soda lakes, where Na^+^ concentrations are elevated (Banciu and Sorokin, 2013; Sorokin et al., 2015). The osmoprotectants accumulate intracellularly to restore the osmotic differential between the environment and the cell (Oren, 1999). The choline-glycine betaine transporter was found to be common among seven of the eight isolate genomes (**Figure 5**) and protects from osmotic stress (Landfald and Strøm, 1986). Smith et al. (1988) found that glycine-betaine cannot be used for biosynthesis and is only synthesized as an osmoprotectant, a direct product of environmental stress response. Accordingly, our experimental design appears to have selected for organisms capable of tolerating the energetic constraints and stressors that would characterize an alkaline, saline, and low free CO_2_ ocean as predicted on Enceladus.

### 4.4 Implications for life detection on Enceladus

The results of this study suggest that metabolically flexible, stress-tolerant heterotrophs capable of exploiting simple organic compounds may be able to survive in putative Enceladus ocean conditions. This has important implications for life detection strategies, which have traditionally focused on identifying canonical metabolic pathways such as autotrophic carbon fixation, particularly methanogenesis. In contrast, our findings indicate that biosignatures associated with metabolic versatility and stress response may be informative targets.

This hypothesis could be further tested through comparative genomics and controlled growth experiments that expose isolates from diverse environments (e.g., soils, marine sediments, and alkaline systems) to Enceladus-relevant stressors, enabling direct comparison of stress-response gene content and expression across ecological contexts. In our dataset, the widespread presence of the F-type H^+^/Na^+^ ATPase suggests that sodium-based bioenergetics may be a conserved strategy under high-pH conditions. As these systems are evolutionarily ancient and central to cellular energy conservation, their breakdown products, in conjunction with measurements of monovalent ion availability, may could serve as biosignatures reflecting adaptation to alkaline ocean chemistry.

In addition, osmotic stress responses may leave detectable molecular fingerprints. The degradation of choline and related osmolytes proceeds through a well-characterized demethylation pathway, producing intermediates including betaine aldehyde, glycine betaine, dimethylglycine, monomethylglycine, glycine, serine, and pyruvate, many of which fall below the molecular size range of complex organics already detected in Enceladus plume particles (Smith et al., 1988; Postberg et al. 2018; Khawaja et al., 2025). The predicted low temperatures, low radiation flux, and largely anoxic conditions on Enceladus may further enhance preservation of such small, labile compounds, increasing their potential as detectable biosignatures of microbial stress response (Davila and Eigenbrode, 2024). These observations suggest that future life detection efforts may benefit from targeting not only bulk organic complexity, but also specific, low-molecular-weight metabolites indicative of physiological adaptation to environmental stress.

## 5. CONCLUSIONS

This microbial enrichment medium informed by Enceladus geochemical data represents one plausible ocean composition, and it can be refined as future missions (e.g., Europa Clipper, JUICE) and re-analyses of Cassini data further constrain the chemistry of Enceladus’ subsurface ocean. Although we supplied H_2_/CO_2_ and acetate to approximate energy sources predicted for Enceladus, these conditions and substrates were selected to reflect plausible environmental chemistry rather than to target specific metabolic pathways.

Within this geochemically bounded medium, we observed the growth of heterotrophs capable of utilizing acetate and exhibiting relatively low carbon requirements, yet these organisms had the genetic potential to metabolize a broad range of carbon substrates. Despite uniformly using acetate and sustaining low carbon demands, the isolate genomes show they can utilize structurally diverse carbon compounds, including short-chain fatty acids, hydrocarbons, and even aromatic substrates. This metabolic flexibility suggests that, even under conditions constrained by Enceladus-like chemistry, some microorganisms capable of surviving may be generalists with broad substrate utilization profiles.

This study highlights the value of using geochemical models to inform the design of microbial growth media and of leveraging terrestrial analog environments as sources of inocula. Although the microorganisms isolated here are not necessarily the most likely candidates for life in Enceladus’ ocean, our results show that members of diverse microbial taxa can survive and grow under conditions predicted for this extraterrestrial environment. Equally important, we provide the medium formulation (**Tables 2 and S1**) as a resource for the broader community to continue this inquiry of microbial habitability in Enceladus-like systems. Future studies could expand upon this work by testing alternative medium formulations, inocula, incubation durations, or cultivation approaches, such as continuous flow chemostats.

## DATA AVAILABILITY

Sequence data is available at NCBI under BioProject accession number PRJNA1457887. Raw genome sequence reads for isolates (*Marinobacter shengliensis*, *Marinobacter vinifirmis, Alishewanella agri*) are available on the Sequence Read Archive (SRA) under accession numbers **SRX33113272, SRX33113271,** and **SRX33113270.** Raw sequence reads for the two mixed cultures (‘metagenomes’) used for binning MAGs are available on SRA under accession numbers **SAMN58702999** and **SAMN58703000.** MAGs from these sequences two metagenomes are available on SRA under accession numbers **SAMN57600512-SAMN57600516.** 16S rRNA genes sequences are available on NCBI’s GenBank with accession numbers **PZ388331-PZ388346**.

## ACKNOWLEDGEMENTS

This work was supported by NASA grants #80NSSC19K1427 to CRG, JAH, JSS, PRG, and ELS, #80NSSC22K1631 to JAH, NSF grant #2023192 to GSS and JAH, National Oceanic and Atmospheric Administration NA19OAR4320072 to JAH, NSF Grant #1921361 to JSS, and the Lewis and Clark Fund for Exploration and Field Research to SE. The field work in Oregon and California was supported by the WHOI Institution funds from philanthropy to the Geodynamics program. TZ was supported by the Massachusetts Department of Education’s STEM Starter Academy through Cape Cod Community College. JAH was also supported by the *Stanley W. Watson Chair for Excellence in Oceanograph*y, while CRG further acknowledges other generous WHOI supporters.

We also thank the captains and crews of the *R/V* Kilo Moana (cruise #KM-2214) and *R/V* Falkor (too) (cruise # FKt230303), as well as the submersible teams for *ROV JASON* and *ROV SuBastian* for their support in acquiring the marine samples used in inoculations. We also acknowledge essential field support from David Butterfield, Bayleigh Benner, Susan Lang, Karen Lloyd, Marc Fontánez-Ortiz, Maddie Peterson, Brandi Kiel Reese, Elizabeth Trembath-Reichert, and C. Geoff Wheat.

## AUTHOR CONTRIBUTIONS

**S.M.E.** Conceptualization, Methodology, Formal Analysis, Investigation, Data Curation, Writing - Original Draft, Writing - Review & Editing, Visualization. **T.E.** Conceptualization, Methodology, Software, Formal Analysis, Investigation, Data Curation, Writing - Original Draft, Writing - Review & Editing, Visualization. **K.W.**, **V.N.**, **L.H.**, **S.M.** Resources, Writing - Review & Editing. **T.Z.**, **A.P.**, **M.S.** Validation, Investigation, Writing - Review & Editing**. E.S.** Writing - Review & Editing, Supervision. **P.G.** Conceptualization, Writing - Review & Editing, Supervision. **C.G.** Writing - Review & Editing, Project Administration, Funding Acquisition. **F.K.** Writing - Review & Editing, Supervision. **J.S.** and **J.A.H.** Conceptualization, Investigation, Resources, Writing - Original Draft, Writing - Review & Editing, Supervision, Project Administration, Funding Acquisition.

## SUPPLEMENTARY INFORMATION

**Table S.1.** Details of Medium 042a preparation

**Table S.2.** Expanded isolate information

**Table S.3.** All pangenome gene clusters

**Table S.4.** Selected pangenome gene clusters

**Table S.5.** Diverse carbon sources used by the isolates

## Footnotes

1 The mathematical center of n-dimensional data clusters, calculated by finding the datapoints in a large cluster that are the shortest distance between the largest number of neighbors

## REFERENCES

1. Adams D (1979) The Hitchhiker’s Guide to the Galaxy. London: Pan Books.

2. Alain K and Querellou J (2009) Cultivating the uncultured: Limits, advances, and future challenges. Extremophiles 13(4): 583–594.

3. Alazard D, Dukan S, Urios A, Verhé F, Bouabida N, Morel F, et al. (2003) *Desulfovibrio hydrothermalis* sp. nov., a novel sulfate-reducing bacterium isolated from hydrothermal vents. International Journal of Systematic and Evolutionary Microbiology 53(1): 173–178.

4. Alneberg J, Bjarnason BS, de Bruijn I, Schirmer M, Quick J, Ijaz UZ, Loman NJ and Quince C (2013) CONCOCT: Clustering contigs on coverage and composition. *arXiv* 1312.4038.

5. Altschul SF, Gish W, Miller W, Myers EW and Lipman DJ (1990) Basic local alignment search tool. Journal of Molecular Biology 215(3): 403–410.

6. Arkin AP, Cottingham RW, Henry CS, Harris NL, Stevens RL, Maslov S, Dehal P, Ware D, Perez F, Canon S, et al. (2018) KBase: The United States Department of Energy Systems Biology Knowledgebase. Nature Biotechnology 36: 566–569. DOI: 10.1038/nbt.4163.

7. Aroney ST, Newell RJ, Nissen JN, Camargo AP, Tyson GW and Woodcroft BJ (2025) CoverM: Read alignment statistics for metagenomics. Bioinformatics 41(4): btaf147.

8. Banciu HL and Sorokin DY (2013) Adaptation in haloalkaliphiles and natronophilic bacteria. In: Seckbach J, Oren A and Stan-Lotter H (eds) Polyextremophiles: Life Under Multiple Forms of Stress. Dordrecht: Springer Netherlands, pp. 121–178.

9. Bankevich A, Nurk S, Antipov D, Gurevich AA, Dvorkin M, Kulikov AS, Lesin VM, Nikolenko SI, Pham S, Prjibelski AD, et al. (2012) SPAdes: A new genome assembly algorithm and its applications to single-cell sequencing. Journal of Computational Biology 19(4): 455–477. DOI: 10.1089/cmb.2012.0021.

10. Baumberger T, Antriasian AM, Butterfield DA, Lu G, Lilley MD and Cathalot C (2023) Gas composition of three new hydrothermal vent sites discovered along the Mid-Atlantic Ridge between 20 and 25° N. AGU Fall Meeting Abstracts 2023: OS13A–05.

11. Bolger AM, Lohse M and Usadel B (2014) Trimmomatic: A flexible trimmer for Illumina sequence data. Bioinformatics 30: 2114–2120. DOI: 10.1093/bioinformatics/btu170.

12. Braun A, Spona-Friedl M, Avramov M, Elsner M, Baltar F, Reinthaler T, Herndl GJ and Griebler C (2021) Reviews and syntheses: Heterotrophic fixation of inorganic carbon – Significant but invisible flux in environmental carbon cycling. Biogeosciences 18: 3689–3700.

13. Bremer E and Krämer R (2019) Responses of microorganisms to osmotic stress. Annual Review of Microbiology 73(1): 313–334.

14. Charlou J-L, Donval J-P, Fouquet Y, Jean-Baptiste P and Holm N (2002) Geochemistry of high H_2_ and CH_4_ vent fluids issuing from ultramafic rocks at the Rainbow hydrothermal field (36°14′N, MAR). Chemical Geology 191: 345–359.

15. Chaumeil P-A, Mussig AJ, Hugenholtz P and Parks DH (2019) GTDB-Tk: A toolkit to classify genomes with the Genome Taxonomy Database. Bioinformatics 36(7): 1925–1927. DOI: 10.1093/bioinformatics/btz848.

16. Daval D, Choblet G, Sotin C and Guyot F (2022) Theoretical considerations on the characteristic timescales of hydrogen generation by serpentinization reactions on Enceladus. Journal of Geophysical Research: Planets 127(2): e2021JE006995.

17. Davila AF and Eigenbrode JL (2024) Enceladus: Astrobiology revisited. Journal of Geophysical Research: Biogeosciences 129(5): e2023JG007677.

18. Delmont TO and Eren AM (2018) Linking pangenomes and metagenomes: The *Prochlorococcus* metapangenome. PeerJ 6: e4320. DOI: 10.7717/peerj.4320.

19. DeLong EF (1992) Archaea in coastal marine environments. Proceedings of the National Academy of Sciences of the United States of America 89(12): 5685–5689.

20. Dilworth MJ and Glenn AR (2007) Problems of adverse pH and bacterial strategies to combat it. In: Novartis Foundation Symposium 221: Bacterial Responses to pH. Chichester: John Wiley & Sons, pp. 4–18.

21. Dougherty MK, Buratti BJ, Seidelmann PK and Spencer JR (2018) Enceladus as an active world: History and discovery. In: Dougherty MK, Esposito LW and Krimigis SM (eds) Enceladus and the Icy Moons of Saturn. Tucson, AZ: University of Arizona Press, pp. 3–25.

22. Eddy SR (2011) Accelerated profile HMM searches. PLoS Computational Biology 7(10): e1002195.

23. Edwards U, Rogall T, Blöcker H, Emde M and Böttger EC (1989) Isolation and direct complete nucleotide determination of entire genes: Characterization of a gene coding for 16S ribosomal RNA. Nucleic Acids Research 17(19): 7843–7853.

24. Eickenbusch P, Takai K, Sissmann O, Suzuki S, Menzies C, Sakai S, Sansjofre P, Tasumi E, Bernasconi SM, Glombitza C and Jørgensen BB (2019) Origin of short-chain organic acids in serpentinite mud volcanoes of the Mariana convergent margin. Frontiers in Microbiology 10: 1729.

25. Elkassas SM, Serres MH, Richardson D, Zhilina TN and Huber JA (2024) Draft genome sequence of *Methanocalculus natronophilus* sp. strain Z-7105T, an alkaliphilic, methanogenic archaeon isolated from a soda lake. Microbiology Resource Announcements 13(7): e00350–24.

26. Ely TD, Leong JM, Canovas PA and Shock EL (2023) Huge variation in H₂ generation during seawater alteration of ultramafic rocks. *Geochemistry, Geophysics*, Geosystems 24: e2022GC010658. DOI: 10.1029/2022GC010658.

27. Eren AM, Kiefl E, Shaiber A, et al. (2021) Community-led, integrated, reproducible multi-omics with anvi’o. Nature Microbiology 6: 3–6. DOI: 10.1038/s41564-020-00834-3.

28. Fifer LM, Catling DC and Toner JD (2022) Chemical fractionation modeling of plumes indicates a gas-rich, moderately alkaline Enceladus ocean. The Planetary Science Journal 3(8): 191.

29. Fryer P, Wheat CG, Williams T, Kelley C, Johnson K, Ryan J, Kurz W, Shervais J, Albers E, Bekins B and Debret B (2020) Mariana serpentinite mud volcanism exhumes subducted seamount materials: Implications for the origin of life. *Philosophical Transactions of the Royal Society A: Mathematical*, Physical and Engineering Sciences 378(2165): 20180425.

30. Galkiewicz JP and Kellogg CA (2008) Cross-kingdom amplification using bacteria-specific primers: Complications for studies of coral microbial ecology. Applied and Environmental Microbiology 74(24): 7828–7831. DOI: 10.1128/AEM.01303-08.

31. Glein CR and Baross JA (2015) The pH of Enceladus’ ocean. Geochimica et Cosmochimica Acta 162: 202–219.

32. Glein CR, Postberg F and Vance SD (2018) The geochemistry of Enceladus: Composition and controls. In: Dougherty MK, Esposito LW and Krimigis SM (eds) Enceladus and the Icy Moons of Saturn. Tucson, AZ: University of Arizona Press, pp. 39–56.

33. Glein CR and Waite JH (2020) The carbonate geochemistry of Enceladus’ ocean. Geophysical Research Letters 47(3): e2019GL085885.

34. Glein CR and Truong N (2025) Phosphates reveal high pH ocean water on Enceladus. Icarus 436: 116717.

35. Gurevich A, Saveliev V, Vyahhi N and Tesler G (2013) QUAST: Quality assessment tool for genome assemblies. Bioinformatics 29(8): 1072–1075. DOI: 10.1093/bioinformatics/btt086.

36. Hand KP, Chyba CF, Priscu JC, Carlson RW and Nealson KH (2009) Astrobiology and the potential for life on Europa. Icarus 200(1): 589–604. DOI: 10.1016/j.icarus.2008.10.011.

37. Harper GD (1984) The Josephine ophiolite, northwestern California. Geological Society of America Bulletin 95(9): 1009–1026.

38. Hilgert U, McKay S, Khalfan M, Williams J, Ghiban C and Micklos D (2014) DNA Subway: Making genome analysis egalitarian. In: Proceedings of the 2014 Annual Conference on Extreme Science and Engineering Discovery Environment. New York, NY: Association for Computing Machinery, pp. 1–3.

39. Horikoshi K (1999) Alkaliphiles: Some applications of their products for biotechnology. Microbiology and Molecular Biology Reviews 63(4): 735–750.

40. Horikoshi K (2021) Alkaliphiles. Boca Raton, FL: Routledge.

41. Hsu H-W, Postberg F, Sekine Y, Shibuya T, Kempf S, Horányi M, Juhász A, Altobelli N, Suzuki K, Masaki Y and Kuwatani T (2015) Ongoing hydrothermal activities within Enceladus. Nature 519(7542): 207–210.

42. Huber JA, Resing JA, Lu GS, Roe KK, Barrett PL, McAllister S, Baumberger T, Beeson JW, Antriasian AM, Merle SG and Walker SL (2024) Multi-vehicle discovery of three new hydrothermal vent fields on the Mid-Atlantic Ridge. In: Proceedings of the Ocean Sciences Meeting 2024, New Orleans, LA, 18–23 February 2024. American Geophysical Union, Abstract DS34A–0389.

43. Hügler M and Sievert SM (2011) Beyond the Calvin cycle: Autotrophic carbon fixation in the ocean. Annual Review of Marine Science 3: 261–289.

44. Hyatt D, Chen G-L, Locascio PF, Land ML, Larimer FW and Hauser LJ (2010) Prodigal: Prokaryotic gene recognition and translation initiation site identification. BMC Bioinformatics 11: 119. DOI: 10.1186/1471-2105-11-119.

45. Jain C, Rodriguez-R LM, Phillippy AM, Konstantinidis KT and Aluru S (2018) High throughput ANI analysis of 90K prokaryotic genomes reveals clear species boundaries. Nature Communications 9(1): 5114. DOI: 10.1038/s41467-018-07641-9.

46. Kang DD, Li F, Kirton E, Thomas A, Egan R, An H and Wang Z (2019) MetaBAT 2: An adaptive binning algorithm for robust and efficient genome reconstruction from metagenome assemblies. PeerJ 7: e7359.

47. Khawaja N, Postberg F and O’Sullivan TR (2025) Detection of organic compounds in freshly ejected ice grains from Enceladus’s ocean. Nature Astronomy. DOI: 10.1038/s41550-025-02655-y.

48. Kim M, Oh H-S, Park S-C and Chun J (2014) Towards a taxonomic coherence between average nucleotide identity and 16S rRNA gene sequence similarity for species demarcation of prokaryotes. International Journal of Systematic and Evolutionary Microbiology 64(Pt 2): 346–351.

49. Kitada M and Horikoshi K (1978) Sodium ion-stimulated α-[1-14C]aminoisobutyric acid uptake in alkalophilic *Bacillus* species. Journal of Bacteriology 131(7): 977–984.

50. Klein F, Bach W and McCollom TM (2013) Compositional controls on hydrogen generation during serpentinization of ultramafic rocks. Lithos 178: 55–69.

51. Koser SA (1948) Growth factors for microorganisms. Annual Review of Microbiology 2(1): 121–142.

52. Landfald B and Strøm AR (1986) Choline-glycine betaine pathway confers a high level of osmotic tolerance in *Escherichia coli*. Journal of Bacteriology 165(3): 849–855.

53. Letunic I and Bork P (2007) Interactive Tree Of Life (iTOL): An online tool for phylogenetic tree display and annotation. Bioinformatics 23(1): 127–128.

54. Li H (2013) Aligning sequence reads, clone sequences, and assembly contigs with BWA-MEM. arXiv 1303.3997.

55. Li H, Handsaker B, Wysoker A, Fennell T, Ruan J, Homer N, Marth G, Abecasis G, Durbin R and 1000 Genome Project Data Processing Subgroup (2009) The sequence alignment/map format and SAMtools. Bioinformatics 25(16): 2078–2079.

56. Lo C-C and Chain PS (2014) Rapid evaluation and quality control of next-generation sequencing data with FaQCs. BMC Bioinformatics 15: 366. DOI: 10.1186/s12859-014-0366-2.

57. Maden BEH (2000) Tetrahydrofolate and tetrahydromethanopterin compared: Functionally distinct carriers in C1 metabolism. Biochemical Journal 350(3): 609–629.

58. Manna V, Balestra C, Banchi E, et al. (2025) High contribution of dark dissolved inorganic carbon uptake to microbial carbon cycling in a shallow Mediterranean basin. Ocean Microbiology 1: 2. DOI: 10.1186/s44375-025-00002-0.

59. Mariner RH, Presser TS, Evans WC and Pringle MK (1990) Discharge rates of fluid and heat by thermal springs of the Cascade Range, Washington, Oregon, and northern California. Journal of Geophysical Research: Solid Earth 95(B12): 19517–19531.

60. McCollom TM and Seewald JS (2007) Abiotic synthesis of organic compounds in deep-sea hydrothermal environments. Chemical Reviews 107(2): 382–401.

61. McCollom TM and Bach W (2009) Thermodynamic constraints on hydrogen generation during serpentinization of ultramafic rocks. Geochimica et Cosmochimica Acta 73(3): 856–875.

62. Merino N, Aronson HS, Bojanova DP, Feyhl-Buska J, Wong ML, Zhang S and Giovannelli D (2019) Living at the extremes: Extremophiles and the limits of life in a planetary context. Frontiers in Microbiology 10: 780601.

63. Montoya L, Celis LB, Razo-Flores E and Alpuche-Solís ÁG (2012) Distribution of CO₂ fixation and acetate mineralization pathways in microorganisms from extremophilic anaerobic biotopes. Extremophiles 16(6): 805–817.

64. Mulkidjanian AY, Galperin MY, Makarova KS, Wolf YI and Koonin EV (2008) Evolutionary primacy of sodium bioenergetics. Biology Direct 3: 13.

65. Nakagawa S and Takai K (2008) Deep-sea vent chemoautotrophs: Diversity, biochemistry and ecological significance. FEMS Microbiology Ecology 65(1): 1–14.

66. Oren A (1999) Bioenergetic aspects of halophilism. Microbiology and Molecular Biology Reviews 63(2): 334–348.

67. Pang KD, Voge CC, Rhoads JW and Ajello JM (1984) The E ring of Saturn and satellite Enceladus. Journal of Geophysical Research: Solid Earth 89(B11): 9459–9470.

68. Parks DH, Imelfort M, Skennerton CT, Hugenholtz P and Tyson GW (2015) CheckM: Assessing the quality of microbial genomes recovered from isolates, single cells, and metagenomes. Genome Research 25(7): 1043–1055. DOI: 10.1101/gr.186072.114.

69. Porco CC, Helfenstein P, Thomas PC, Ingersoll AP, Wisdom J, West R, Neukum G, Denk T, Wagner R, Roatsch T and Kieffer S (2006) Cassini observes the active south pole of Enceladus. Science 311(5766): 1393–1401.

70. Postberg F, Kempf S, Schmidt J, Brilliantov N, Beinsen A, Abel B, Buck U and Srama R (2009) Sodium salts in E-ring ice grains from an ocean below the surface of Enceladus. Nature 459(7250): 1098–1101.

71. Postberg F, Schmidt J, Hillier J, Kempf S and Srama R (2011) A salt-water reservoir as the source of a compositionally stratified plume on Enceladus. Nature 474(7353): 620–622.

72. Postberg F, Khawaja N, Abel B, Choblet G, Glein CR, Gudipati MS, Henderson BL, Hsu H-W, Kempf S, Klenner F and Moragas-Klostermeyer G (2018) Macromolecular organic compounds from the depths of Enceladus. Nature 558(7711): 564–568.

73. Postberg F, Sekine Y, Klenner F, et al. (2023) Detection of phosphates originating from Enceladus’s ocean. Nature 618: 489–493. DOI: 10.1038/s41586-023-05987-9.

74. Rodionov DA, Vitreschak AG, Mironov AA and Gelfand MS (2002) Comparative genomics of thiamin biosynthesis in procaryotes. Journal of Biological Chemistry 277(50): 48949–48959.

75. Randolph-Flagg NG, Ely T, Som SM, Shock EL, German CR and Hoehler TM (2023) Phosphate availability and implications for life on ocean worlds. Nature Communications 14(1): 2388.

76. Raux E, Schubert HL and Warren MJ (2000) Biosynthesis of cobalamin (vitamin B12): A bacterial conundrum. Cellular and Molecular Life Sciences 57(13–14): 1880–1893.

77. Rempfert KR, Miller HM, Bompard N, Nothaft D, Matter JM, Kelemen P, Fierer N and Templeton AS (2017) Geological and geochemical controls on subsurface microbial life in the Samail Ophiolite, Oman. Frontiers in Microbiology 8: 56.

78. Richter M and Rosselló-Móra R (2009) Shifting the genomic gold standard for the prokaryotic species definition. Proceedings of the National Academy of Sciences of the United States of America 106(45): 19126–19131.

79. Saitou N and Nei M (1987) The neighbor-joining method: A new method for reconstructing phylogenetic trees. Molecular Biology and Evolution 4(4): 406–425.

80. Sañudo-Wilhelmy SA, Gómez-Consarnau L, Suffridge C and Webb EA (2014) The role of B vitamins in marine biogeochemistry. Annual Review of Marine Science 6(1): 339–367.

81. Seewald JS (2017) Detecting molecular hydrogen on Enceladus. Science 356(6334): 132–133.

82. Sekine Y, Shibuya T, Postberg F, Hsu H-W, Suzuki K, Masaki Y, Kuwatani T, Mori M, Hong PK, Yoshizaki M and Tachibana S (2015) High-temperature water-rock interactions and hydrothermal environments in the chondrite-like core of Enceladus. Nature Communications 6: 8604.

83. Seth EC and Taga ME (2014) Nutrient cross-feeding in the microbial world. Frontiers in Microbiology 5: 350.

84. Shaffer M, Borton MA, McGivern BB, Zayed AA, La Rosa SL, Solden LM, Liu P, Narrowe AB, Rodríguez-Ramos J, Bolduc B, Gazitúa MC, Daly RA, Smith GJ, Vik DR, Pope PB, Sullivan MB, Roux S and Wrighton KC (2020) DRAM for distilling microbial metabolism to automate the curation of microbiome function. Nucleic Acids Research 48(15): 8883–8900. DOI: 10.1093/nar/gkaa621.

85. Silantyev SA, Buikin AI, Gurenko AA, Chugaev AV, Shabykova VV, Tskhovrebova AR, Beltenev VE and Bich AS (2024) Geochemical signature of basalts of the MAR Rift Valley at 20°31′N: Origin conditions of the anomalous volcanic center of Puy des Folles in the axial zone of the Mid-Atlantic Ridge. Geochemistry International 62(11): 1111–1122.

86. Smith LT, Pocard JA, Bernard T and Le Rudulier D (1988) Osmotic control of glycine betaine biosynthesis and degradation in *Rhizobium meliloti*. Journal of Bacteriology 170(7): 3142–3149.

87. Sorokin DY (2005) Is there a limit for high-pH life? International Journal of Systematic and Evolutionary Microbiology 55(4): 1405–1406.

88. Sorokin DY, Banciu HL and Muyzer G (2015) Functional microbiology of soda lakes. Current Opinion in Microbiology 25: 88–96.

89. Spahn F, Schmidt J, Albers N, Hörning M, Makuch M, Seiss M, Kempf S, Srama R, Dikarev V, Helfert S, Moragas-Klostermeyer G, Krivov AV, Sremcevic M, Tuzzolino AJ, Economou T and Grün E (2006) Cassini dust measurements at Enceladus and implications for the origin of the E ring. Science 311(5766): 1416–1418. DOI: 10.1126/science.1121375.

90. Stackebrandt E and Liesack W (1993) Nucleic acids and classification. In: Goodfellow M and O’Donnell AG (eds) Handbook of New Bacterial Systematics. London: Academic Press, pp. 152–189.

91. Stackebrandt J and Ebers J (2006) Taxonomic parameters revisited: Tarnished gold standards. Microbiology Today 33: 152–155.

92. Stecher G, Tamura K and Kumar S (2020) Molecular evolutionary genetics analysis (MEGA) for macOS. Molecular Biology and Evolution 37(4): 1237–1239.

93. Stern JC, Graham HV, Burcar B, Martin ES, Noell A, Hand K, et al. (2025) A comprehensive framework for assessing terrestrial analogue field sites for ocean worlds. Journal of Geophysical Research: Planets 130: e2024JE008803. DOI: 10.1029/2024JE008803.

94. Takai K, Moyer CL, Miyazaki M, Nogi Y, Hirayama H, Nealson KH and Horikoshi K (2005) *Marinobacter alkaliphilus* sp. nov., a novel alkaliphilic bacterium isolated from subseafloor alkaline serpentine mud from Ocean Drilling Program Site 1200 at South Chamorro Seamount, Mariana Forearc. Extremophiles 9(1): 17–27.

95. Tamura K, Nei M and Kumar S (2004) Prospects for inferring very large phylogenies by using the neighbor-joining method. Proceedings of the National Academy of Sciences of the United States of America 101(30): 11030–11035.

96. Tamura K, Stecher G and Kumar S (2021) MEGA11: Molecular evolutionary genetics analysis version 11. Molecular Biology and Evolution 38(7): 3022–3027. DOI: 10.1093/molbev/msab120.

97. Taylor SJ and Krejca JJ (2006) *A Biological Assessment of Caves in Lava Beds National Monument*. Technical Report BIOD 2006-06. National Park Service.

98. Trembath-Reichert E, Shah Walter SR, Ortiz MA, Carter PD, Girguis PR and Huber JA (2021) Multiple carbon incorporation strategies support microbial survival in cold subseafloor crustal fluids. Science Advances 7(18): eabg0153.

99. Tucholke BE and Schouten H (1988) Kane fracture zone. Marine Geophysical Researches 10(1): 1–39.

100. Waite JH Jr, Combi MR, Ip W-H, Cravens TE, McNutt RL Jr, Kasprzak W, Yelle R, Luhmann J, Niemann H, Gell D and Magee B (2006) Cassini ion and neutral mass spectrometer: Enceladus plume composition and structure. Science 311(5766): 1419–1422.

101. Waite JH Jr, Lewis WS, Magee BA, Lunine JI, McKinnon WB, Glein CR, Mousis O, Young DT, Brockwell T, Westlake J and Nguyen MJ (2009) Liquid water on Enceladus from observations of ammonia and 40Ar in the plume. Nature 460(7254): 487–490.

102. Waite JH, Glein CR, Perryman RS, Teolis BD, Magee BA, Miller G, Grimes J, Perry ME, Miller KE, Bouquet A and Lunine JI (2017) Cassini finds molecular hydrogen in the Enceladus plume: Evidence for hydrothermal processes. Science 356(6334): 155–159.

103. Wolery TW and Jarek RL (2003) *Software User’s Manual: EQ3/*6, Version 8.0. Livermore, CA: Lawrence Livermore National Laboratory.

104. Wolin HL and Naylor HB (1957) Basic nutritional requirements of *Micrococcus lysodeikticus*. Journal of Bacteriology 74(2): 163–167.

105. Wolin EA, Wolin MJ and Wolfe RS (1963) Formation of methane by bacterial extracts. Journal of Biological Chemistry 238(8): 2882–2886.

106. Yuen G, Blair N, Des Marais DJ and Chang S (1984) Carbon isotope composition of low molecular weight hydrocarbons and monocarboxylic acids from the Murchison meteorite. Nature 307(5948): 252–254.

107. Zhang Y and Gladyshev VN (2008) Trends in selenium utilization in the marine microbial world. Biochemical and Biophysical Research Communications 371(1): 1–5.

108. ZoBell CE (1941) Studies on marine bacteria. I. The cultural requirements of heterotrophic aerobes. Journal of Marine Research 4(1): 42–75.

109. Zolotov MY (2007) An oceanic composition on early and today’s Enceladus. Geophysical Research Letters 34(23): L23203.

